# Traumatic brain injury causes retention of long introns in metabolic genes via regulation of intronic Histone 3 lysine 36 methylation levels in the sub-acute phase of injury

**DOI:** 10.1101/077214

**Authors:** Arko Sen, Wen Qu, Oluwademi Okikiolu Nuga, Jenney Liu, Rayanne Burl, Katherine Gurdziel, Roger Pique-Regi, Maik Hüttemann, Douglas. M. Ruden

## Abstract

Traumatic brain injury (TBI) can cause persistent pathological alteration of neurons. This may lead to cognitive dysfunctions, depression, and even increased susceptibility to life threatening diseases, such as epilepsy and Alzheimer’s Disease. To investigate the underlying genetic and molecular basis of TBI, Wasserman and colleagues developed an inexpensive and reproducible model for simulating TBI in *Drosophila melanogaster* (Fruit Fly). Using a modified version of this high impact trauma (HIT) device, we subjected w^1118^ fruit flies to mild closed head trauma. To determine the transcriptomic changes that contribute to survival post TBI, we collected fly heads from the survivors at 2 time points; 4 hours and 24 hours’ post-trauma. Mild TBI had limited impact on the steady state RNA levels but showed large perturbations in alternative splicing (AS) 24 hours’ post-trauma. Classification of these AS changes showed selective retention (RI) of long introns (>81bps), with a mean size of ~3000bps. Some of these RI genes also showed a significant reduction in transcript abundance and were specifically enriched in genes involved in mitochondrial metabolism. The RI are enriched in ACACACA motifs known to bind to Smooth (SM), an hnRNPL class of splicing factor. Mutating SM (sm^4^/Df) resulted in reversal of RI observed 24 hours’ post-trauma, and in some cases, elimination of basal levels of RI in long introns. This observation suggests that SM is critical regulator of RI and that this process is enhanced by TBI. Interestingly, chromatin immunoprecipitation followed by deep sequencing (ChIP-seq) for histone 3 lysine 36 trimethylation (H3K36me3) revealed increased levels of this histone modification in retained introns post-trauma. Consistent with this observation, mutations in lysine specific demethylase 4A (KDM4A), which de-methylate H3K36me3, increased RI in many of the same long introns affected by TBI. Additionally, higher H3K36me3 levels are observed around intronic SM-binding motifs post-trauma, suggesting interaction between H3K36me3 and SM binding to intronic splicing suppressor sites might be responsible for increasing RI of metabolic genes as a novel mechanism to improve survival after TBI.

## Introduction

Traumatic brain injury (TBI) is a complex pathological condition associated with high mortality rates (Unterharnscheidt 1995; Ling et al. 2015). TBI can be divided into 2 major categories; severe TBI or concussive brain injury and mild TBI (MTBI) or sub-concussive brain injury (Broglio et al. 2012). Severe TBI is typically characterized by dramatic changes in neuronal homeostasis including rapid influx of calcium, efflux of potassium, release of neurotransmitters, and widespread neuronal death (Giza and Hovda 2014). In contrast, sub-concussive trauma has been suggested to cause multifocal microscopic axonal damage and micro-hemorrhages (McKee and Robinson 2014; Daneshvar et al. 2015). A wide spectrum of neurological abnormalities due to severe TBI or MTBI is thought to culminate into larger heterogeneous group of neurological disorders broadly classified as post-concussion syndrome (PCS) (Hall et al. 2005). It may lead to progressive neurodegeneration and increased susceptibility to diseases such as Alzheimer’s disease, Parkinson’s disease, or motor neuron disease (Ling et al. 2015).

Several factors influence the pathology of TBI. Sullivan et al, 2013 demonstrated that sagittal injury is associated with significantly more axonal damage and behavioral changes compared to axial injury in gyrencephalic piglets (Sullivan et al. 2013). This is consistent with primate models, where head rotation and coronal acceleration affects the diffusion pattern of axonal injury and the severity of coma (Gennarelli et al. 1982; Browne et al. 2011). Besides the primary factors such as location and strength of the injury, the genotype of an individual may also contribute to an increase in susceptibility to neuronal dysfunction post-TBI. For example, inheritance of the Apoliprotien E (Apo ε4) allele is associated with increased deposition of Aβ protein, increased severity of chronic neuronal diseases and increased risk of Alzheimer’s disease (AD) post-TBI (Dardiotis et al. 2010). Similarly, specific mutations in α-synuclein are associated with increased risk of Parkinson’s disease after TBI (Acosta et al. 2015).

Considering the heterogeneity of factors that can influence pathological outcomes post-TBI, there is an immediate need to develop animal models to study the molecular mechanisms associated with TBI. To this end, Wassarman’s lab developed a model of closed head trauma (CHI) in Drosophila melanogaster (Fruit Fly) (Katzenberger et al. 2013). The authors demonstrated that it is possible to reproduce defining characteristics typical to CHI in humans such as temporary incapacitation, ataxia, and neurodegeneration. For example, Drosophila showed a reduction in lifespan, dependent upon the number of strikes, similar to reports from epidemiological studies in humans. The authors also demonstrated appearance of vacuolar spot in the Drosophila brain post-trauma indicative of neurodegeneration. Furthermore, the number and size of these spots were dependent on age of the flies and the number of strikes. Other laboratories also reported an association between age, the number of primary TBI incidences and progressive neurodegeneration in mice models of TBI (Loane et al. 2014).

Alternative splicing is a major process that contributes to modulation of transcriptome diversity in various pathological conditions. In a recent study, Song et al, 2015, demonstrated that in Drosophila, RNA repair and splicing pathways can influence downstream splicing of the stress responsive gene, Xbp1(Song et al. 2015). This alteration in RNA splicing affects the ability of neurons to regenerate from axotomy or acute neuronal damage. Furthermore, studies in mouse models have demonstrated TBI can cause increased expression of RNA Binding Motif 5 and RNA Binding Motif 10 proteins that coincide with neurodegeneration in the hippocampus (Jackson et al. 2015). However, most studies have been limited because they looked at gene-specific changes in alternative splicing in response to neuronal damage rather than global changes. Therefore, in this study we attempt to define the scope and diversity of splicing changes at a whole genome level after TBI. Additionally, we provide some strong evidence for potential regulatory mechanisms that may control these alternative splicing changes.

## Results

### 1. Increase in number of strikes result in progressive decrease in long-term survival

We simulated CHI is Drosophila using a similar experimental set-up, as described by the Wassarman lab (Katzenberger et al. 2013). Briefly, 0-5 days-old w^1118^ flies were collected in sturdy vials and TBI was inflicted using an in-house HIT (high impact trauma) device. Instead of using a 90° spring deflection, we used a 45° spring deflection to attenuate the impact (Fig 1A). We performed survival estimation, after 1 or 2 strikes, using 20 w^1118^ flies per vial and repeated the experiment 5 times for male and female flies, separately. The objective was to determine if we could replicate the strike dependent decrease in survival, as observed by Katzenberger et al, 2013 (Katzenberger et al. 2013). The results as illustrated in Fig. 1B demonstrated that the HIT model of TBI in Drosophila is reproducible and reliable. As noted previously, the female flies seemed to be marginally more resistant to TBI compared to the males. Additionally, most flies which survived past 24 hours post-TBI survived for >10days.

**Figure 1.**
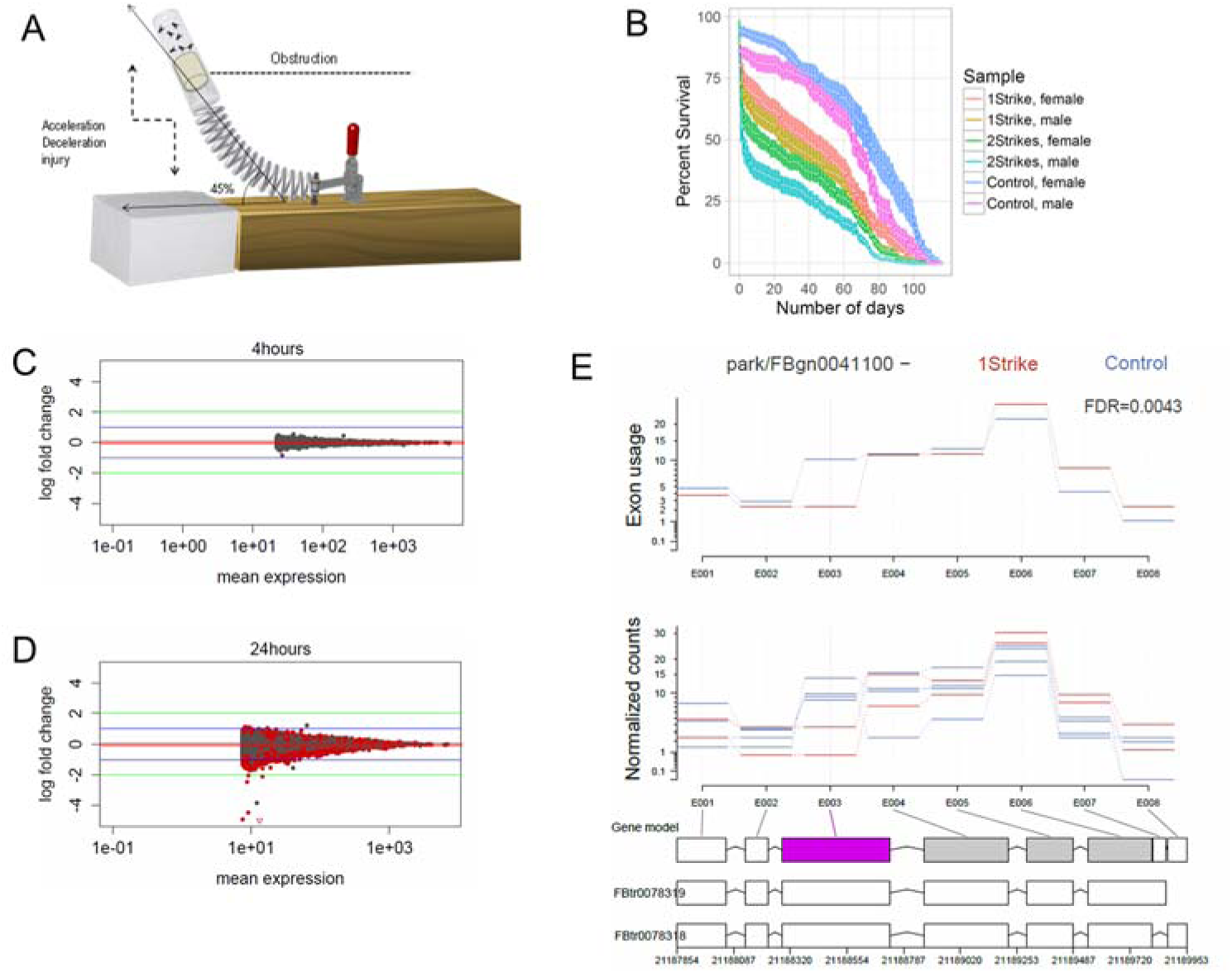
Characterization of exon usage profile of TBI flies. A) Illustration of the modified high impact trauma device (HIT). B) Survival curve (mean ± standard error) of Drosophila subjected to either 1 strike or 2 strike. We used using 20 w^1118^ flies per vial and repeated the experiment 5 times for male and female flies. C) D) MA plots differentially used exons for TBI heads collected 4 hours post-TBI and 24 hours post-TBI. Red dots indicate the genes associated with FDR corrected p-value ≤ 0.1 or 10%. Green horizontal line indicate a logFC ≥ ±2 and blue horizontal line indicate a logFC ≥ ±1.E) FBgn0041100(park) shows changes in exon usage post-TBI. The first panel shows the estimated changes in exon usage. The second panel shows normalized count per exonic part. The bottom panel shows the gene model used for estimating exon usage. In pink are the exons which show a significant change (FDR ≤ 0.1 or 10%) in exon usage. Only exon 3 showed logFC =-1.05 fold change in exon usage.

### 2. Exploratory analysis of RNA-sequencing data revealed mild-changes in expression and exon usage profile

We were interested in studying transcriptional changes associated with MTBI or sub-concussive injury; therefore, for the remainder of the study, we used the 1-strike model for MTBI. Post-TBI we collected the heads of the survivors at 2 time points; 4 hours after TBI and 24 hours after TBI and characterized the expression profiles using RNA-sequencing (Love et al. 2014). Principle component analysis (PCA) of library-size normalized read counts (by exons per gene) demonstrated large variability in between replicate control and TBI samples (Suppl. Fig 1B and C). PCA showed that the majority of the differences in the RNA profiles between control heads (4 and 24 hours non-TBI heads) and heads collected 24 hours post-TBI was attributed to injury (57%), followed by sex of the flies (27%) and finally, 16% by other unknown factors (Suppl. Fig 1C). In the case of heads collected 4 hours after TBI, a majority of the variability relative to the control heads (4 and 24 hours non-TBI heads) were because of sex specific differences (40%), followed by the Injury (35%) and 25% was contributed by other unknown variables (Suppl. Fig 1B). For gene-specific differential expression analyses, we used an additive linear model with sex and time as covariates and used the non-TBI heads collected at 24 hours and 4 hours as controls (see Methods). Our analysis revealed that the number of differentially expressed (DE) genes (TBI vs control) at an FDR corrected p-value cut-off of 0.1 was much greater at 24 hours (N= 1698) (Suppl. Fig 1F) compared to 4 hours (N=145) (Suppl. Fig 1D). To further refine our data, we introduced an additional filter of log2FC≥ ±2 fold. We found only 1 gene, LSP2, that showed a significant change in gene expression for heads collected 4 hours post-TBI and 1 gene, Jon25Bi, which showed expression changes 24 hours after traumatic brain injury. Decreasing the log2FC cut-off from ±2fold to ±1fold result in discovery of an additional 222 DE genes in heads collected 24 hours after TBI and 17 DE genes in heads collected 4hrs after TBI. Our result suggests that attenuated traumatic brain injury (or MTBI) causes only mild changes in steady-state RNA levels of genes and most of these changes are only observed at 24 hours post-TBI.

We performed gene ontology (GO) analysis of the 223 genes showing mild change in expression using DAVID (https://david.ncifcrf.gov/) and filtered the output by FDR corrected p-values ≤ 0.1. Our results showed the top 10 GO classes belonging to regulation of mitochondrial function, translation and mitotic spindle formation associated processes (Suppl. Fig 2). GO analysis was not feasible for the small number of differentially expressed genes detected 4 hours post-TBI.

### 3. Increase in the number of exons showing differential exon usage 24 hours post-TBI

RNA sequencing data can be used to infer alternative splicing profiles of the genome. We inferred the relative abundance of splice isoforms by comparing the exon usage profiles between the control and the experimental (or TBI) samples (Li et al. 2015). Exon usage is the ratio of normalized read counts of the target exon to the normalized read count of any other exon of the transcript (see Methods). Comparing the exon usage profiles between control heads and heads collected at 4 hours post-TBI showed no difference in the exon usage profile (Fig. 1C). However, comparison of exon usage profiles between control heads and heads collected at 24 hours post-TBI showed a large number of changes in exon usage (N=5984/77026) at an FDR corrected p-value cut-off of 0.1 or 10% (Fig. 1D). We further filtered the differentially used exons by a log2FC cut-off of ±2 fold. This reduced our list of differentially used exons to 6 exons. Reducing the log2FC cut-off from ±2 fold to ±1 fold (i.e., from 4-fold to 2-fold changes) increased the number of differentially used exons detected to 124 (Fig. 1D). An example of gene specific change in exon usage in the *park* gene is shown in Figure 1G.

### 4. Characterization of alternative splicing events 24 hours post-TBI using MISO

Our analysis until now exclusively relied on reads mapping to exonic regions and exon-intron junctions. However, studies have demonstrated that inclusion of reads mapping to intronic regions can significantly improve detection of alternative splicing changes, especially intron retention (RI). Therefore, we re-analyzed our RNA-sequencing data using the commonly used splicing analysis software MISO (Mixture of Isoforms)(Katz et al. 2010). Using exon-centric analysis in the software, MISO, we looked for differential alternative splicing events that occurred at 24 hours post-TBI. We filtered our differential splicing events using difference in <u>P</u>ercent <u>S</u>pliced <u>I</u>n (∆PSI) ≥ ±0.05 or ±5% and Bayes Factor ≥ 10 (see MISO manual). We found that the vast majority of the events detected are RI (retained intron; Fig 2A and B). We further determined the linear correlation coefficient (r^2^) between the ∆PSI observed for males and females for all sex-independent differential splicing events (N=578). We observed a ~47% correlation between the ∆PSI values for males and females for RI events (N=458) (Fig 2B) suggesting significant variation between males and females. However, these events are characterized by a positive (+) ∆PSI 24 hours post-TBI independent of sex (Fig 2A and 2B).

**Figure 2.**
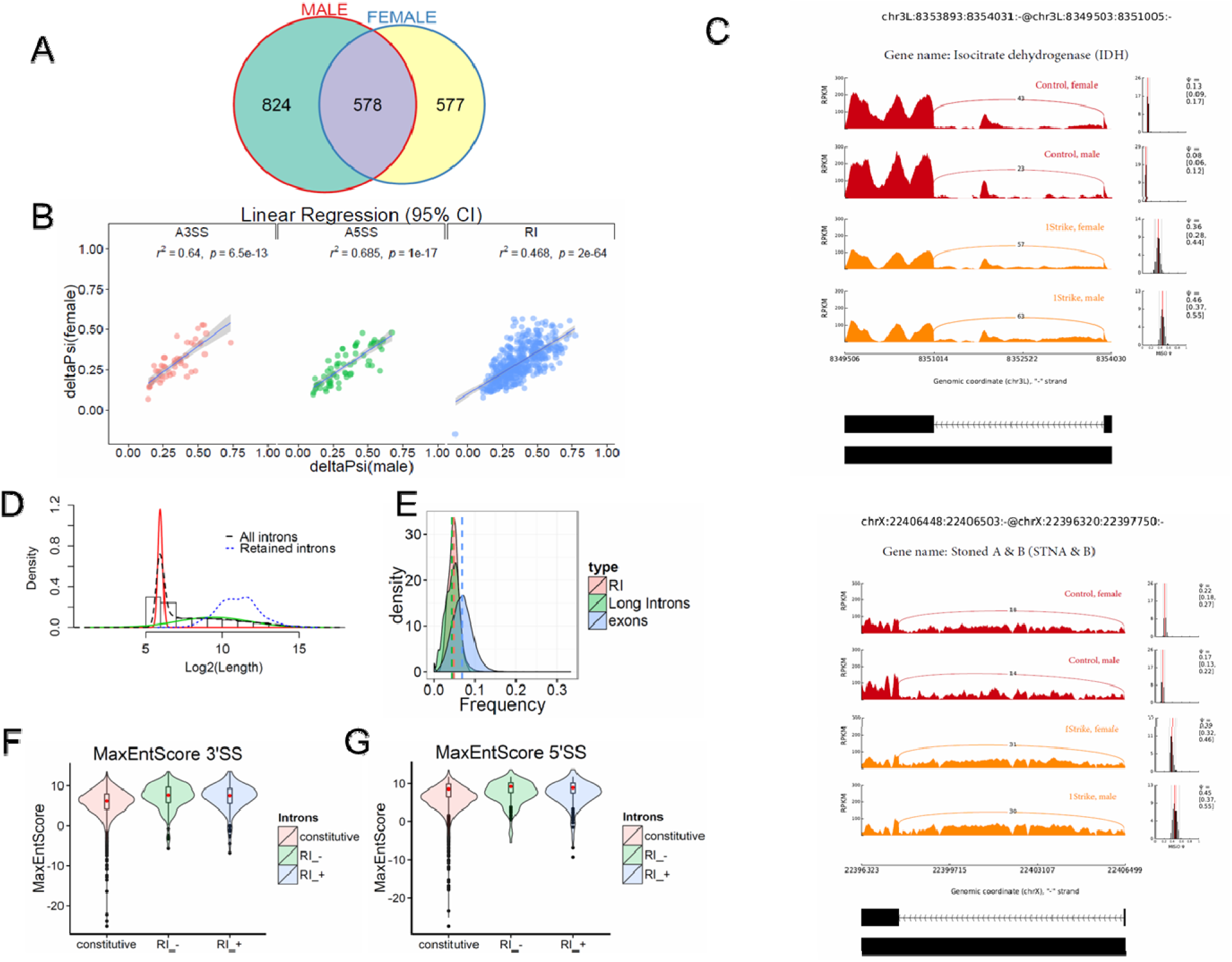
Characterization of alternative splicing events in Drosophila model of TBI. A) Overlaps between statistically significant differential alternative splicing events detected using MISO for heads collected 24 hours post-TBI for males and females. B) Linear regression between ∆PSI or deltaPSI for differential splicing events detected for heads collected 24 hours post-TBI for males and females. C) *sashimi plot* showing intron retention event in *Isocitrate dehydrogenase* (IDH)(panel 1) and stoned A/B (panel 2). The data is represented as Reads Per Kilo-base per Million. D) Graphical representation of *Gaussian mixture model* to determine the natural cut-off for calling long introns. Black dotted line represents the log_2_(intron length) for all introns in the genome. Blue dotted line represents the intron length of retained introns detected for heads collected 24 hours post-TBI for males and females. The red and green line represents the normal distributions fitted using *Gaussian mixture model*. E) Distribution of GC-content for retained introns (RI), long introns and exons. Dotted lines are the median GC-content. Distribution of Maximum entropy scores (MaxEntScores) for F) 3’SS and G) 5’SS for constitutive introns, retained introns (+ strand) (RI_+), and retained introns (- strand) (RI_-).

Gene ontology analysis (see Methods) for all genes (N= 374) mapping to differential RI events showed enrichment (FDR≤0.1) of genes mapping to synaptic vesicle transport and lipid particles. As we collected RNA from whole heads, we anticipated intron retention to affect genes associated with synaptic processes. Manually curating genes containing RI revealed several interesting genes that may explain the loss of long-term fitness in TBI flies. These genes include metabolic genes such as Isocitrate Dehydrogenase (IDH) (Fig 2C, Panel I) and Aconitase (ACON; not shown), and synaptic transport proteins such as Stoned A and B (STN A and B) (Fig 2C, Panel II).

RI is affected by a number of characteristics such as intron length, GC content, and splice-site strength. We calculated the natural cut-off for long introns to be >81bps using a Gaussian mixture model (Fig 2D) (Lim et al, 2001). Using the computed natural cut-off, we observed that all (458/458) of the RIs occurred in long introns. Additionally, these introns have a similar mean GC-content compared to any other long intron in the Drosophila genome (Fig 2E). Studies have demonstrated that it is possible to differentiate between true and decoy splice sites using maximum entropy scores (MaxEntScore). Using the model proposed by Yeo et al, 2004 (Yeo and Burge 2004), we calculated the MaxEntScore of 5’SS and 3’SS short sequence motifs for constitutive introns and retained introns. We defined constitutively spliced introns (constitutive) in a stringent manner as introns that are efficiently processed and are not present in any sample. The maximum entropy scores at the 5’SS and 3’SS for retained introns are not significantly different from constitutive introns (Fig 2F and 2G). This suggests that the RI events are not located at bad decoy splice sites but are located at *bona fide* and good splice sites (Fig 2F and 2G).

The consequence of RI could be the targeting of transcripts to nonsense-mediated decay (NMD) through introduction of a pre-mature termination codon (PTC) (Chang et al. 2007; Isken and Maquat 2007). To test this possibility, we calculated the transcripts per million (TPM) for all individual transcripts (N= 1477) which contained at least one predicted RI. Then we performed a Fisher exact test to determine the significant differences in TPM between the control heads and 24 hours post-TBI heads. 223 transcripts out of 1477 tested were significant at an FDR ≤ 0.05, ∆TPM≥ ±20 (Fig 3A).

**Figure 3:**
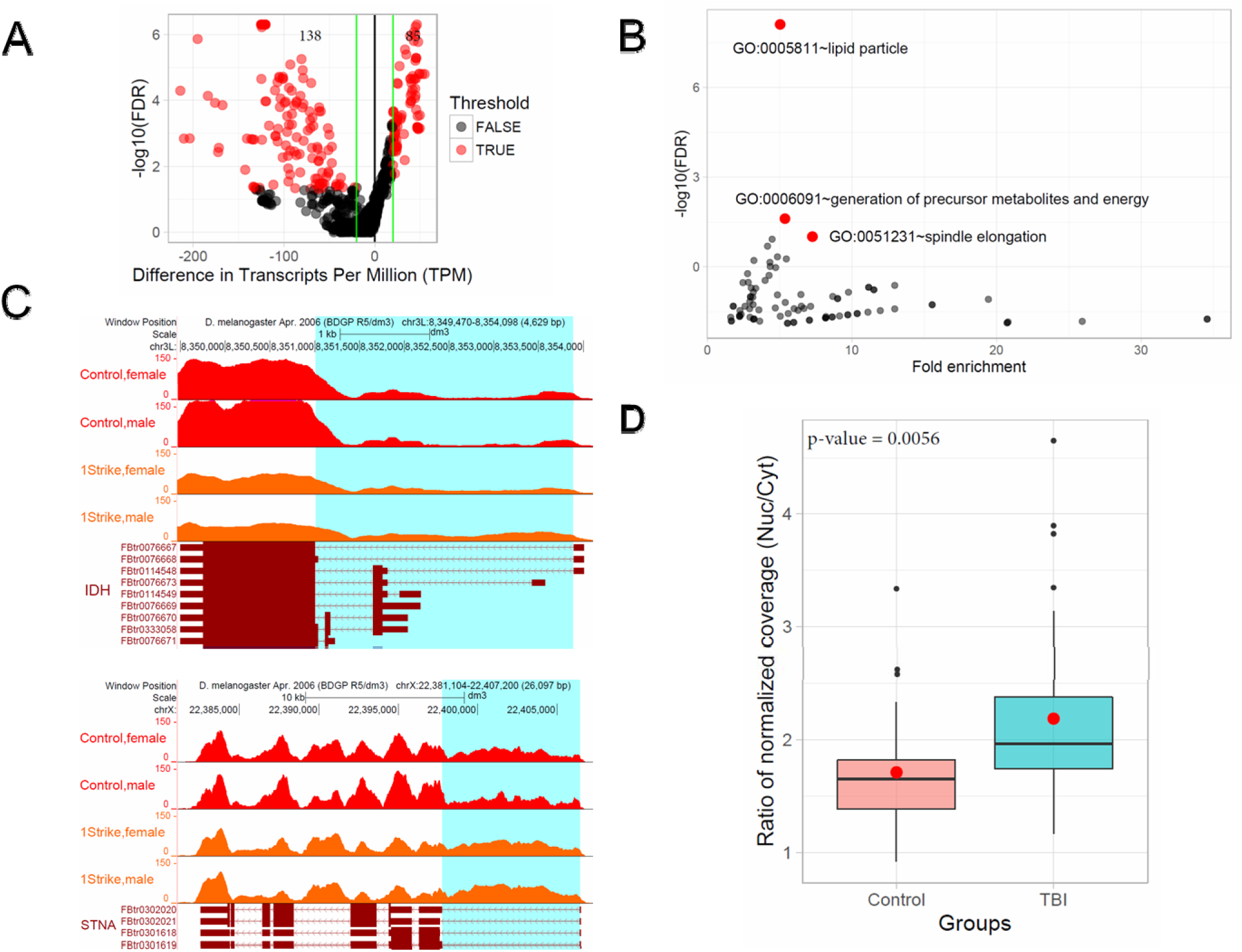
Intron retention causes degradation or nuclear retention of transcripts. A) Difference in Transcript abundances (∆TPM) was plotted against –log10 (FDR) for 24 hors post-TBI samples. Red dots indicate introns showing significant change in PIR at a FDR corrected p-value ≤ 0.05. Green horizontal line indicate a ∆TPM ≥ ±20. 138 transcripts showed decrease in TPM and 85 showed increase B) GO analysis was performed in DAVID using 59 genes. 4209 genes which contained introns detected in both male and female heads were used as control. The fold enrichment and −log_10_ (FDR) was calculated and plotted. Red dots indicate the GO categories significant at FDR ≤ 0.1 C) Normalized coverage in exons and introns for Isocitrate dehydrogenase (IDH) across control and TBI (24 hours post-TBI) samples (panel 1). Normalized coverage in exons and introns for Stoned A and B(STN A and B) across control and TBI (24 hours post-TBI) samples (panel 2). TBI result in RI of the highlighted introns in figure C panel 1 and 2. D) Normalized read coverage was calculated for Nuclear and Cytoplasmic mRNA fraction for control and 24 hours post-TBI samples for genes (N=32) showing increase in transcript abundance on RI post-trauma. Ratio of Nuclear/Cytoplasmic normalized read coverage is plotted on Y axis. Differential testing was done using Welch 2 sample t.test.

Using this Method, we observed that the transcripts separated out into 2 distinct groups. The first group (N=138) of RI transcripts showed a reduction in steady-state RNA levels. Transcripts in this group belonged to a set of 59 genes which were highly enriched (FDR≤0.05) for GO categories such as GO:0005811~lipid particles, GO:0006091~generation of precursor metabolites and energy and GO:0051231~spindle elongation (Fig 3B). An interesting example in the metabolite category is Isocitrate dehydrogenase (IDH) which showed a significant decrease in expression across males and females 24 hours post-TBI (Fig 3C, Panel I).

The second group of RI transcripts (n=85) (Fig 3A and D) showed a paradoxical increase in steady-state RNA levels. One possible explanation for this observation could be that RI causes “nuclear retention” and stabilization of the transcripts. To test this possibility, we collected RNA from the nuclear and cytoplasm fractions from control heads and 24 hours post-TBI heads. For each sample, we calculated the normalized coverage for genes that showed an increase in transcript abundance with RI post-TBI. Then we calculated the ratio of normalized read coverage for nuclear and cytoplasmic RNA fractions. The expectation was that this ratio should be significantly higher for TBI samples. Consistent with our hypothesis, we observed accumulation of transcripts which show increase in transcript abundance on RI in the nuclear fraction 24 hours post-TBI (Fig 3D). In contrast, the RI transcripts that showed a decrease in transcript abundance were primarily cytosolic (data not shown).

Studies have demonstrated that splicing in Drosophila occurs co-transcriptionally. This can be tested directly from the RNA-seq data by calculating the splicing rate (SR) (Wickramasinghe et al. 2015). Furthermore, this study also demonstrated that SR can be used to make qualitative assessments concerning the efficiency of co-transcriptional splicing. Briefly, splicing rate (SR) is the ratio of the normalized-count for the first intron and the normalized-count for the last-intron. SR ≥ 1 indicates post-transcriptional processing while SR < 1 indicates co-transcriptional processing. The SR analysis was only restricted to genes with a single transcript (see Methods) (N=756). These filters were necessary because SR calculations are confounded by potential complexity introduced by other types of alternative splicing events that would give artificially low SRs.

The SR for each sample was divided into *quartiles* and only the first 3 *quartiles* were plotted to control for the effect of outliers. We used the Wilcoxon rank sum test to check for differences in SR, and observed that the median SR for TBI and control flies to be <1. This indicates that co-transcriptional splicing is the predominant mechanism of splicing in Drosophila (Fig. 6A). Additionally, we saw a statistically significant increase (p-value from Wilcoxon rank sum test < 2.2e-16) in SR in post-TBI heads indicating decreased splicing efficiency (Wickramasinghe et al. 2015) (Fig 6A and Suppl. Fig 3A and B).

### 5. Mitochondrial profile in the 24 hours post-TBI

We observed downregulation in the expression of several genes involved in the mitochondrial trichloroacetic acid (TCA) cycle associated with RI in 24 hours post-TBI heads such as Pyruvate kinase (PYK), Enolase(ENO), Aconitase (ACON) and Isocitrate dehydrogenase (IDH). Therefore, we hypothesized that dysregulation of the TCA cycle genes will lead to mitochondrial stress post-TBI. To test this hypothesis, we tested the effect of TBI on mitochondrial cytochrome *c* oxidase (COX), the terminal and rate-limiting enzyme of the electron transport chain (Villani et al. 1998; Kunz et al. 2000). We found that COX activity increased by 22% in the TBI heads compared to controls (Fig. 4A). This suggests that an increased energy demand is likely due to tissue remodeling after trauma, a process that is energy expensive. In fact, analysis of ATP levels indicated a significant 27.2% decrease of cellular ATP in the 24 hours post-TBI heads (Fig. 4B). A drop of cellular ATP and concomitant increase in ADP would further activate COX since this enzyme is allosterically regulated by the ATP/ADP ratio (Arnold and Kadenbach 1999).

**Figure 4:**
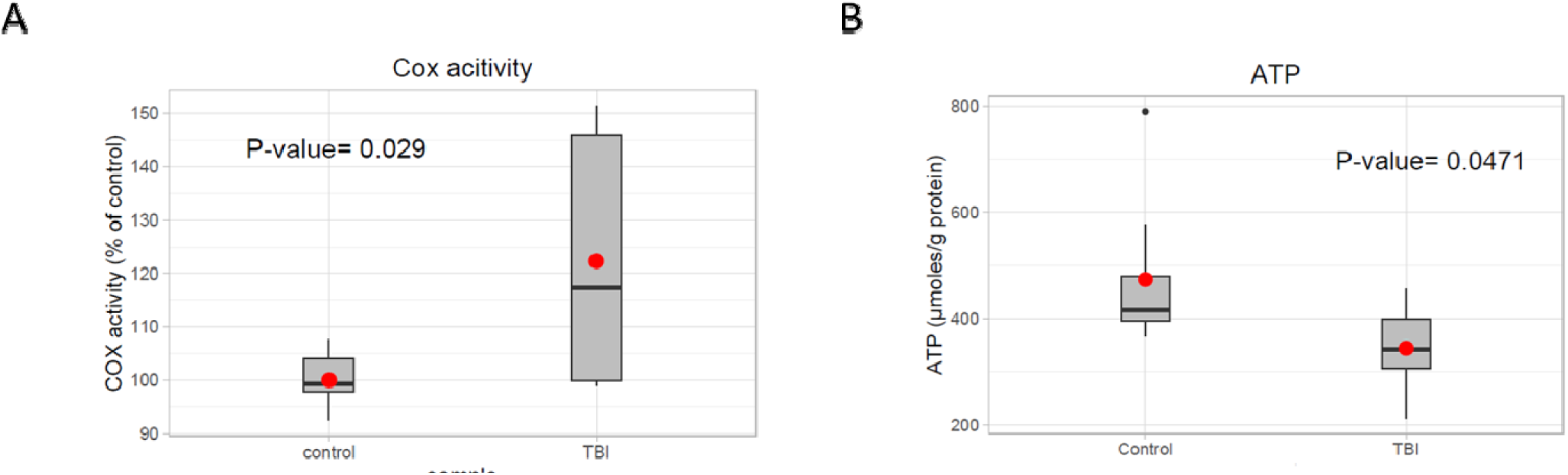
Mitochondrial profile post-TBI. A) Effect of TBI on COX activity in total fly heads (see methods). COX specific activity of solubilized heads was measured using the polarographic method and is reported as % of control. The 24 hours post-TBI heads show an 22% increase in COX activity (p.value = 0.029). B) ATP levels in fly heads were determined using the bioluminescent method and normalized to protein content. Heads collected 24 hours post-TBI shows a significant decrease (p.value= 0.0471) in total ATP concentration (µmol,ATP/mg Protein) compared to the control.

### 6. H3K36me3 is depleted from alternative exons compared to constitutive exons indicating that it may play a regulatory role in splicing

In a previous section, we demonstrated that splicing in Drosophila takes place co-transcriptionally. Therefore, histone modifications might have an important role to play in splicing regulation by interacting with the elongating RNA polymerase. One of the major histone modifications implicated in splicing is H3K36me3 (Luco et al. 2010; Pradeepa et al. 2012; Zhou et al. 2014). The methyl groups on H3K36me3 are removed from the lysine residue by Jumanji-C (JmjC) domain-containing histone demethylases; specifically; KDM4A in Drosophila (Lin et al. 2012; Crona et al. 2013). Therefore, we hypothesized that a reduction of KDM4A protein level will cause significant changes in alternative splicing in a manner that resembles TBI. Homozygous loss of function mutations of KDM4A in Drosophila cause developmental arrest and do not allow survival beyond the early embryonic stage (Tsurumi et al. 2013). Therefore, we attempted to conditionally knockdown KDM4A in adult heads. Unfortunately, we were unable to get an appreciable and stable knockdown of the gene in adult heads using the gene-switch system by feeding adults RU486 to induce elav-Gal4-GS mediated induction of UAS-KDM4A-RNAi (not shown). Instead, since KDM4A has been shown to be well expressed in 3^rd^ instar larvae (Lorbeck et al. 2010), we determined the impact of KDM4A reduction on splicing regulation in 3^rd^ instar larvae (see Methods).

We knocked down KDM4A in 3^rd^ instar larvae using the UAS/GAL4 RNAi system (see Methods) (Suppl. Fig 4A) and performed RNA-sequencing using 100bps paired-end reads. KDM4A activity is associated with transcriptional repression in mammalian cells (Zhang et al. 2005; Huang and Dixit 2011). Consistent with this observation, differential expression analysis comparing dKDM4A-RNAi to controls (w^1118^) showed a much larger number of genes with increases in expression at an FDR corrected p-value ≤ 0.1 (10%) and a log2FC ≥ ±2 in 3^rd^ instar larvae (Suppl. Fig 4B).

Comparing the alternative splicing profile of KDM4A-RNAi with the control (w^1118^) 3^rd^ instar larvae, we found 98 splicing events that are well-correlated between biological replicates (Fig 5A). These 98 splicing events showed a significant change in ∆PSI values (∆PSI ≥ |0.05| or |5%| and Bayes Factor ≥ 10). Among them majority of the events (70/98) was RIs (Fig 5B). These events showed a significant 52% positive correlation between replicates. Notably, 29/70 events were also detected in 24 hours post-TBI heads (Fig 5D), including RIs detected in metabolic regulators that were also observed in the TBI experiments, such as IDH and PYK and genes responsible for synaptic vehicle endocytosis such as STNA and STNB(Fig 5E).

**Figure 5:**
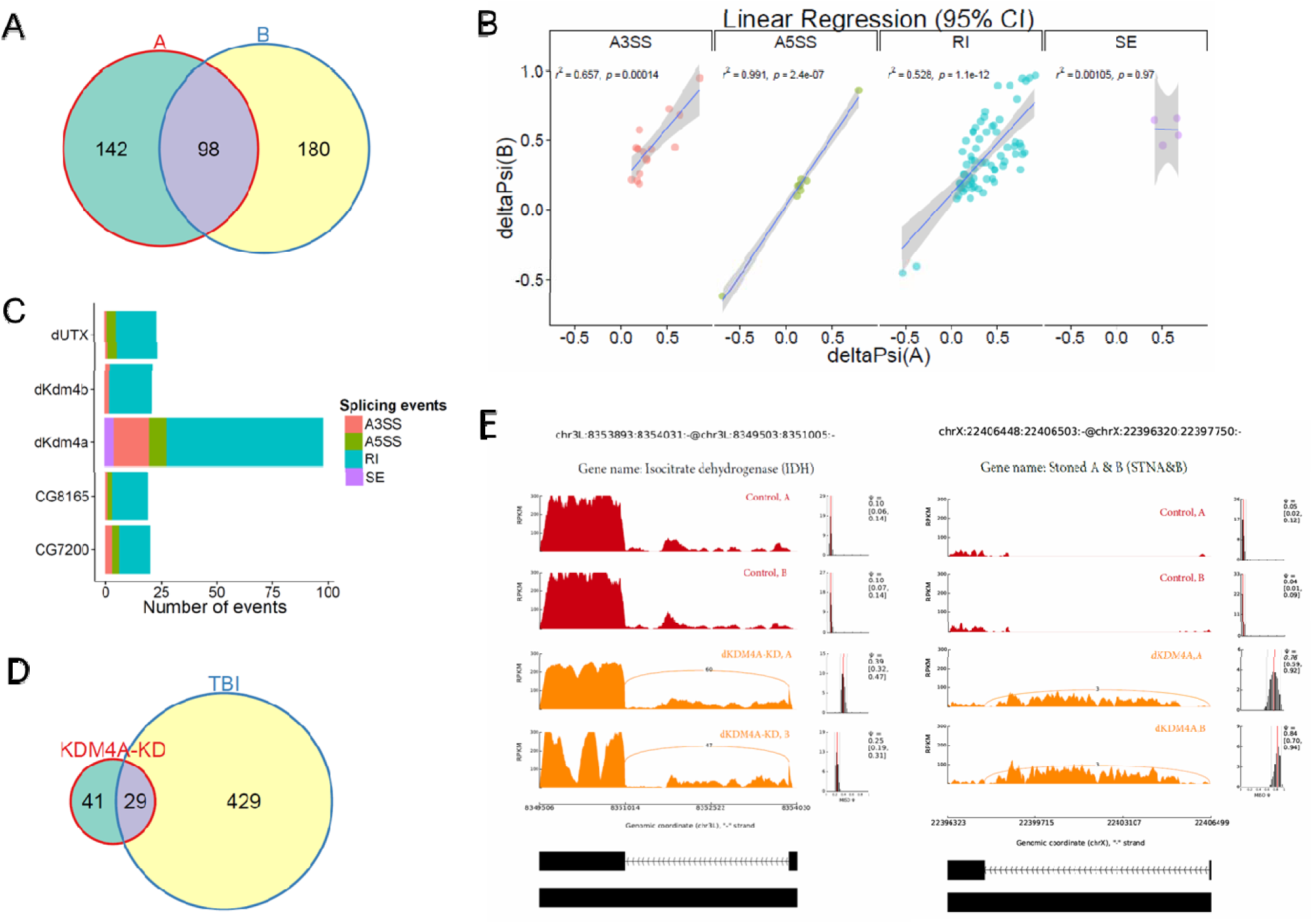
Knockdown of KDM4A in 3rd instar larvae causes Intron retention in Isocitrate dehydrogenase (IDH) A) Overlap between differential alternative splicing events detected in biological replicates. B) Linear regression between ∆PSI or deltaPSI for differential splicing events detected for dmutantM4A-mutant samples across biological replicates (A,B). C) Positive control experiment showing that dmutantM4A-mutant show the biggest changes in alternative splicing compared of other histone demethylases; dUTX(H3K27 demethylase), CG8165 (H3K9me2/me3 demethylase), CG7200 (prospective demethylase containing jmjC domain), dmutantM4B (H3K9 and H3K36 demethylase, lower affinity compared to dmutantM4A). D) Overlaps between statistically significant differential alternative splicing events detected using MISO for heads collected 24 hours post-TBI for males and females and dmutantM4A-mutant in 3rd instar larvae. E) sashimi plot showing intron retention event in Isocitrate dehydrogenase (IDH) (left panel) and Stoned A (STNA) (right panel). The data is represented as Reads Per Kilo-base per Million.

The KDM4A reduction results provided us with initial evidence that H3K36me3 might play an integral role in regulating RIs. To determine if this association is unique for H3K36me3, we performed RNA-sequencing of RNAi lines for the H3K27me3-specific histone demethylase, UTX (Copur and Muller 2013), and the H3K9me2/3-specific demethylases KDM3A (CG8165) (Herz et al. 2014), KDM4B and JmjC-domain containing uncharacterized protein CG7200 in Drosophila pupae. The expectation was that if H3K27me3 and H3K9me2/me3 do not contribute to alternative splicing, reduction of the regulatory histone demethylases will have limited effects on the splicing profiles. Consistent with our expectation, splicing analyses showed only 23 differential splicing events for UTX-RNAi, 19 differential splicing events for KDM3A (CG8165)-RNAi, 21 differential splicing events for KDM4B-RNAi and 20 differential splicing events for CG7200-RNAi (Fig 5C). None of the differential splicing events in any of the other KDM-RNAi samples were detected in the KDM4A-RNAi or 24 hours post-TBI samples. Therefore, our knock-down experiments with Drosophila 3^rd^ instar larvae suggested that H3K36me3 may play a critical role in regulating alternative splicing.

To determine the characteristic profile of H3K36me3 in Drosophila tissues, we used online H3K36me3 chromatin-immunoprecipitation followed by next-generation DNA-sequencing (ChIP-seq) (35 bps single-end) data for 3^rd^ instar larvae (L3) (GSE47248) and heads (GSE47280) from the modENCODE consortium. We called significantly-enriched peaks against the respective input controls within 200bps non-overlapping sliding windows across the entire genome. Then we filtered them using a FDR corrected p-value ≥ 0.05 and log2FC≥ 2. Finally, we calculated the mean RPKM of significant peaks across disjointed and unique exonic and intronic regions of transcripts and plotted their density distributions using R/Bioconductor. Our analysis indicated that exons are generally marked by a higher level of H3K36me3 compared to introns in Drosophila 3^rd^ instar larvae (Fig. 6B) and adult heads (Fig. 6C). This observation was also noted in our H3K36me3 ChIP sequencing (100 bps paired-end) data (Fig 6D).

**Figure 6.**
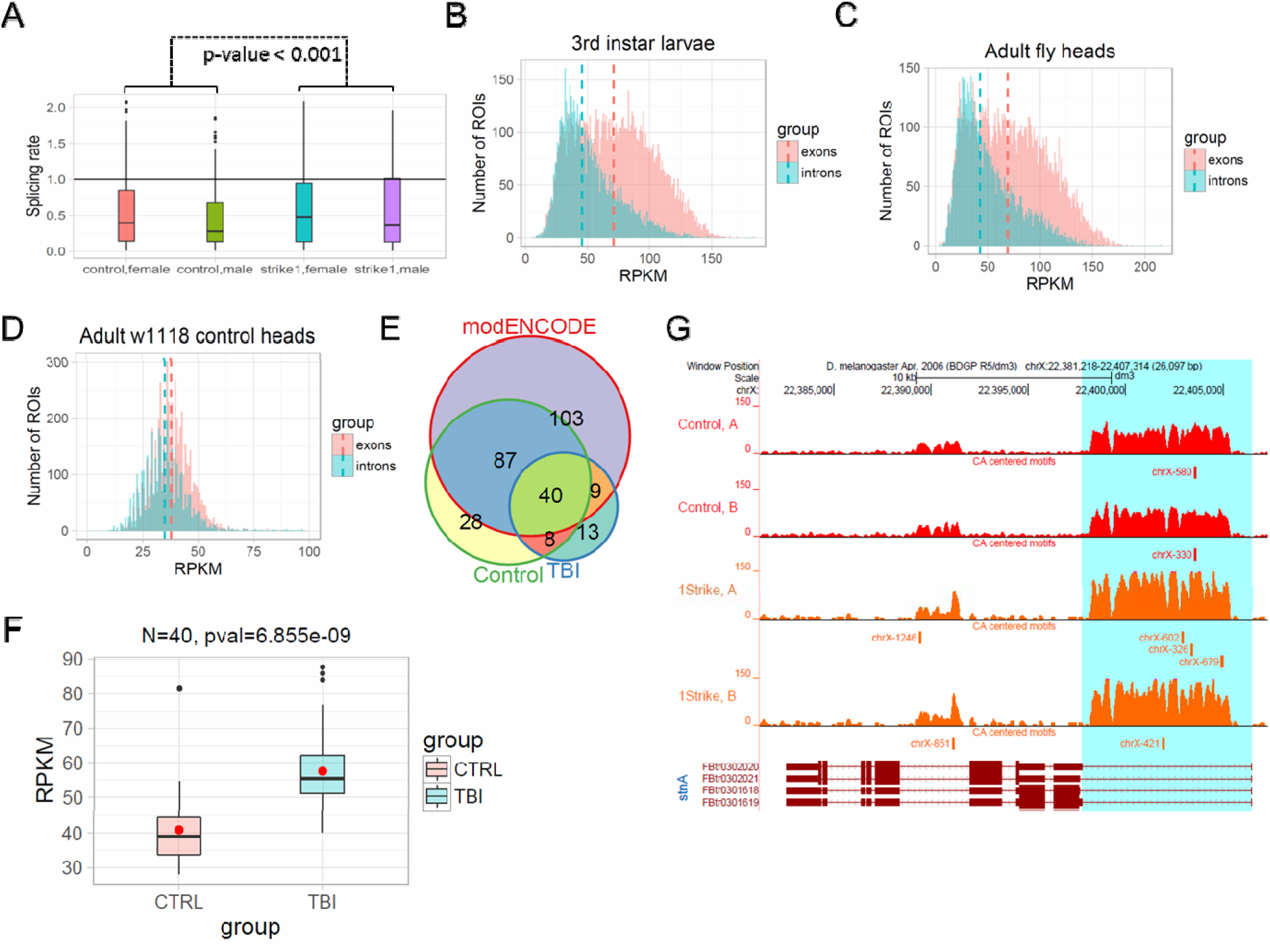
H3K36me3 is an potential regulator of alternative splicing: A) Distribution of Splicing Rate (SR) plotted for control and 1 strike samples (1 strike, 24 hours post-TBI) for Drosophila model of TBI. Splicing rate for all samples were divided into quartiles and only the first 3 quartiles where plotted to control for the effect of outliers. Wilcoxon rank sum test was performed to check for statistically significant difference in SR between control and 1 strike samples; p-value < 0.001. Histogram of mean normalized counts from ModENCODE chip-seq data on introns and exons for B) 3rd instar larvae C) Adult fly heads. The differential tag densities within 200 bp regions was estimated using MEDIPS. Regions which showed a ≥ + 2 fold enrichment at an FDR corrected p-value ≤ 0.05 over input, was considered as significant peaks. The target peaks was overlapped with regions of interest (ROIs). Criteria for selecting ROIs are as follows; introns and exons containing significant peaks was ≥200 bps in length. Reads Per Kilobase per Million (RPKM) for respective ROIs were plotted in R/Bioconductor. D) The differential tag densities within 200 bp windows was estimated using MEDIPS. Regions which showed an enrichment at an FDR corrected p-value ≤ 0.1 over input, was considered as significant peaks to maximize peak calls. H3k36me3 detected for w1118 control fly heads were mapped to exons and introns (≥200bps in length). RPKM distribution and median RPKM for all peaks mapping to exonic and intronic regions were plotted using R/Bioconductor.E) The RI contained significant H3K36me3 peaks for Control, TBI and modENCODE H3k36me3 ChIP data were overlapped. We found 40 RI in common. F) For these 40 RI Average RPKM for all ChIP peaks were estimated for Control and TBI samples. These 40 intron showed an significant increase in H3K36me3 in TBI samples (pval = 6.855e-09, Welch 2 sample T-test). G) D)The RI in STNA shows a general increase in H3K36me3 in TBI compared to w1118 control. We remapped our significant peaks and centered it around CA-rich motifs (ACACACA with 1 mismatch). The bars at the bottom of each track shows the location of the CA-centered significant peaks.

To further our understanding of the H3K36me3 profile and the possible association between RI and TBI, we used the available exon annotation from ENSEMBL to classify constitutive and alternative exons into groups of 3 (i.e., trios). The constitutive exon trio (CE trios) consisted of 3 adjacent CEs. The alternative trio (AE trios) consisted of 1 AE flanked by 2 CEs in a constitutive-alternative-constitutive configuration (see Methods). We observed that the H3K36me3 was depleted from the AE relative to the flanking CE in both 3^rd^ instar larvae and adult heads (Suppl. Fig 6A and B). Moreover, the CE trios showed a completely different pattern compared to AE trios (Suppl. Fig 6A and 5B). One-sided ANOVA showed a significant difference between the alternative and constitutive exons for the alternative trio (Suppl. table 1) (P-adjusted ≤0.1) in 3^rd^ instar larvae and the pattern was well conserved from the 3^rd^ instar larvae to adult heads. This suggests that the relationship between H3K36me3 and AS is well-conserved throughout development. Meta-analysis of mod-ENCODE data and RNA-sequencing of KDM4A-RNAi in 3^rd^ instar larvae and adult fly heads, suggests that H3K36me3 is associated with regulation of alternative splicing post-TBI. This encouraged us to perform H3K36me3 ChIP-seq analysis of Drosophila heads ±TBI to decipher this histones mark’s association with RI.

### 7. H3K36me3 is associated with splicing changes in the 24 hours post-TBI

Based on our reanalysis of modENCODE ChIP-Seq data and KDM4A knockdown studies in 3^rd^ instar larvae, we hypothesized that post-TBI there is an increase in H3K36me3 within the introns in the RI genes in the heads. To confirm this hypothesis, we performed ChIP for H3K36me3 in w^1118^ control and TBI heads and sequenced them using 100bps paired-end reads. The long 100bps paired-end reads are advantageous as they allow better estimation of the immunoprecipitated fragment size distribution than the 36bps reads in the modENCODE dataset (Approx. 176bps for all samples). We called enriched peaks against input controls within 200bps sliding windows using an FDR corrected p-value cut-off of 0.1 (10%) for w^1118^ control and 24 hours post-TBI fly heads (see Methods). Annotation of significant H3K36me3 peaks showed higher enrichment of peaks mapping to intronic and distal intergenic regions in 24 hours post-TBI heads and exonic regions in control (w^1118^) heads. This suggested a general genome-wide shift in the H3K36me3 pattern post-trauma to favor intronic regions (Fig 6E).

Following peak annotation, we calculated the mean-RPKM of H3K36me3 peaks within our RIs (see Methods). The RIs which contained a minimum of 8 reads per samples were considered as represented in our final dataset. We detected 48/458 RIs which contained significant peaks in the control (w^1118^) and 24 hours post-TBI heads. To provide an independent validation of our H3K36me3 ChIP, we also calculated the mean-RPKM of significant peaks for modENCODE H3K36me3 heads (GSE47280) within RIs. Again, only RIs with minimum of 8 reads per sample were selected for further study. 40/458 RIs were represented in control (w^1118^), 24 hours post-TBI heads and modENCODE H3K36me3 ChIP-seq datasets (Fig 6F). For these 40 RIs, the mean-RPKM of H3K36me3 peaks for 24 hours post-TBI heads were significantly higher compared to the control (w^1118^) heads as illustrated in Fig 6G. Normalized coverage plots of H3K36me3 for representative RIs in STNA and RpS4 are illustrated in Supplemental Fig 6C and D. Interestingly, for STNA we observed a base level of RI that correlates with higher H3K36me3 in long introns compared to surrounding exons. In the TBI heads, the H3K36me3 level is higher in the STNA RI, which corresponds to increased inclusion of the long intron (Suppl. Fig 6C and D). Therefore, this suggests that RI might the mechanism for modulating expression of transcripts, especially of certain neuronal transport proteins.

### 8. Discovery of splicing factor binding motifs

The association between H3K36me3 and RI can be further impacted by interaction with RNA binding proteins (RBPs). RBPs can bind conserved sequences within intron and regulate their efficient processing and subsequent removal. De-novo motif enrichment analysis using DREME, within 300 bps windows near the 5’SS and 3’SS of RIs revealed specific enrichment of CA-rich motifs RNA-binding motifs (Table1 and 2). This CA-rich RNA motif binds to Smooth (SM), an hnRNPL (heteronuclear ribonuclear protein-like) homologue (Suppl. Fig. 10), or CNOT4, an RBP protein of unknown function (Ray et al. 2013). We also found other motifs for RBPs such as SHEP (AT-rich motif), ARET (GT-rich motif) and RBP9 (T rich motif) near the 5’SS (Table 1), as well as SHEP (AT-rich motif) and SF1 near the 3’SS (Table 2). Motif analyses were performed using reshuffled 300 sequences as controls (see Methods). We repeated our motif enrichment analysis using randomly selected 300 bps regions from the Drosophila genome(dm3) as controls using the program motifRG in R/Bioconductor (see Methods). Using motifRG, we observed similar enrichment of CA-rich motif within 300 bps of the 5’SS but not the 3’SS in RIs. However, we did not see enrichment of any of the other (non-CA) repeats using motifRG (Suppl. Table 2 and 3).

**Table1:**
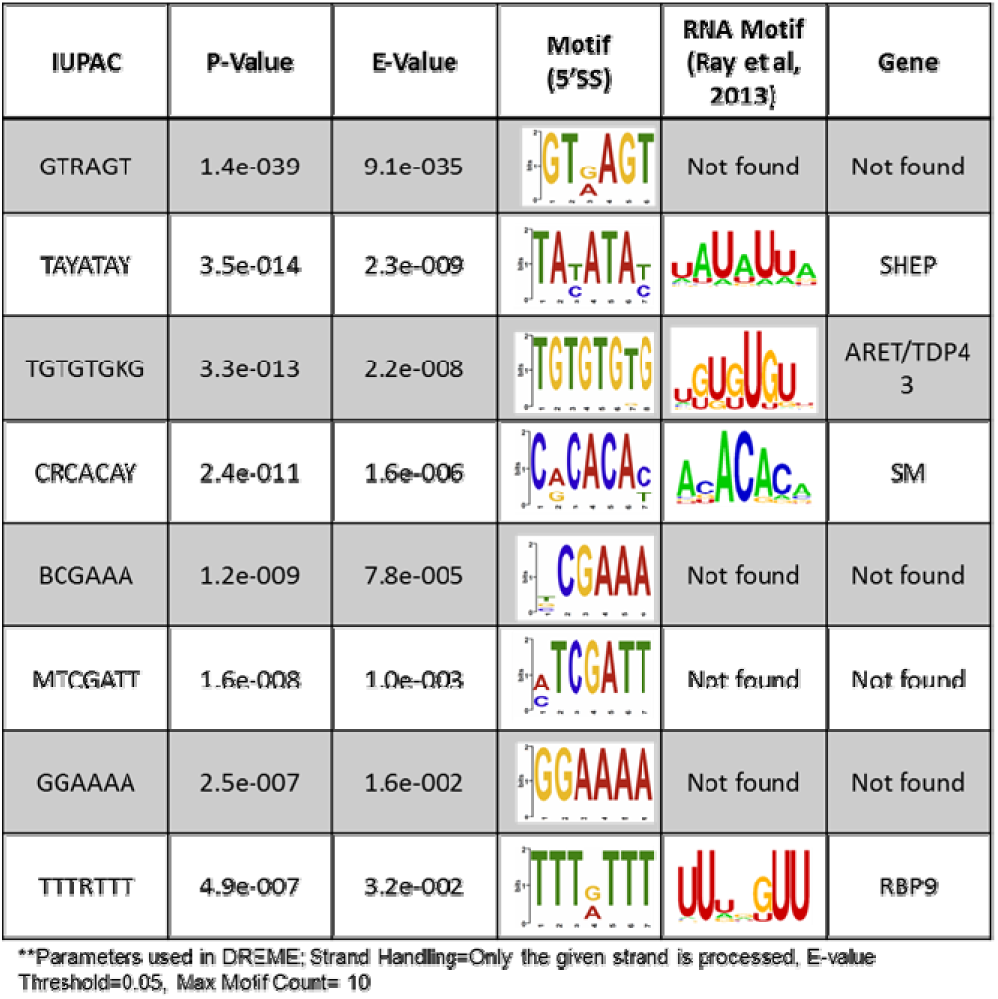
*Motif enrichment analysis for 5’SS*. 299 bps sequences were collected from intronic side of 5’SS for retained intron of length ≥ 600 (N= 421/458). Motif enrichment analysis was run using reshuffled sequences as background (Parameters; Strand Handling=Only the given strand is processed, E-value Threshold=0.05, Max Motif Count= 10).

**Table2:**
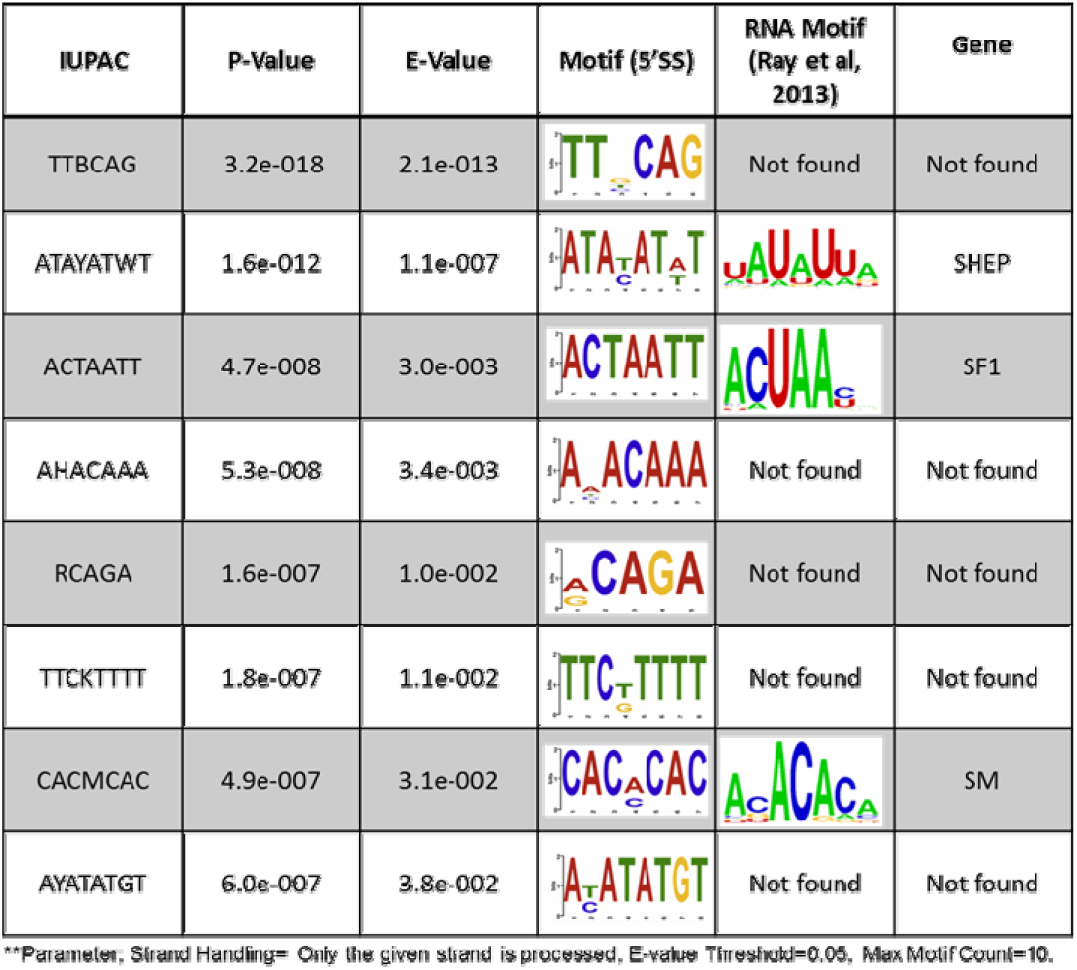
*Motif enrichment analysis for 3’SS*. 299 bps sequences were collected from intronic side of 5’SS for retained intron of length ≥ 600 (N= 421/458). Motif enrichment analysis was run using reshuffled sequences as background (Parameters; Strand Handling=Only the given strand is processed, E-value Threshold=0.05, Max Motif Count= 10).

Downregulation of expression of splicing factors binding to these enriched motifs 24 hours post-TBI might be responsible for RI. However, we observed that the all the detected splicing factor with enriched motifs except SF1 were downregulated in 3^rd^ instar larvae and pupae and showed the higher expression in 0-5 days old heads (Suppl. Figure 5A). Also, they did not show any change steady-state RNA levels 24 hours post-TBI. Furthermore, it also suggested that differential binding rather than altered RNA levels post-TBI might be associated with splicing regulation.

As the SM-binding CA-rich motif is near (within 300bps) the 5’SS and 3’SS of the RI with DREME and near the 5’SS using motifRG, we considered SM to be our strongest candidate for further analysis. The frequency of ACACACA motif allowing 1 mismatch was significantly higher in long introns (≥81 bps) compared to short introns (Suppl. Fig 5B). Therefore, we hypothesized that the presence of CA-rich motifs in long introns might be a defining characteristic of Drosophila long introns that is required for its efficient processing.

To understand the role of SM in regulation of splicing events we obtained the sm4 (SM) mutant stock. Briefly, the sm4 mutant is a mutant of the SM gene and is semi-lethal over a deficiency (Df) (Karpen and Spradling 1992). These sm^4^/Df flies survive to adulthood but are short lived with a median age of about ~30 days (Layalle et al. 2005). After confirming the short-lived phenotypic characteristic of the sm^4^/Df flies (data not shown), we subjected them to a 1 strike and collected the heads 24 hours +/- TBI and performed RNA-seq analyses. We estimated the ∆PSI using the MISO compared to the w^1118^ controls separately for males and females and filtered them using ∆PSI ≥ ±5 and Bayes factor ≥ 10. We only considered the events that overlapped between males and females for further analysis. We found that only 17 RIs detected 24 hours Post-TBI were detected in sm^4^/Df +TBI which meant that sm4/Df mutation rendered the flies non-responsive to TBI (Fig 7A).

**Figure 7:**
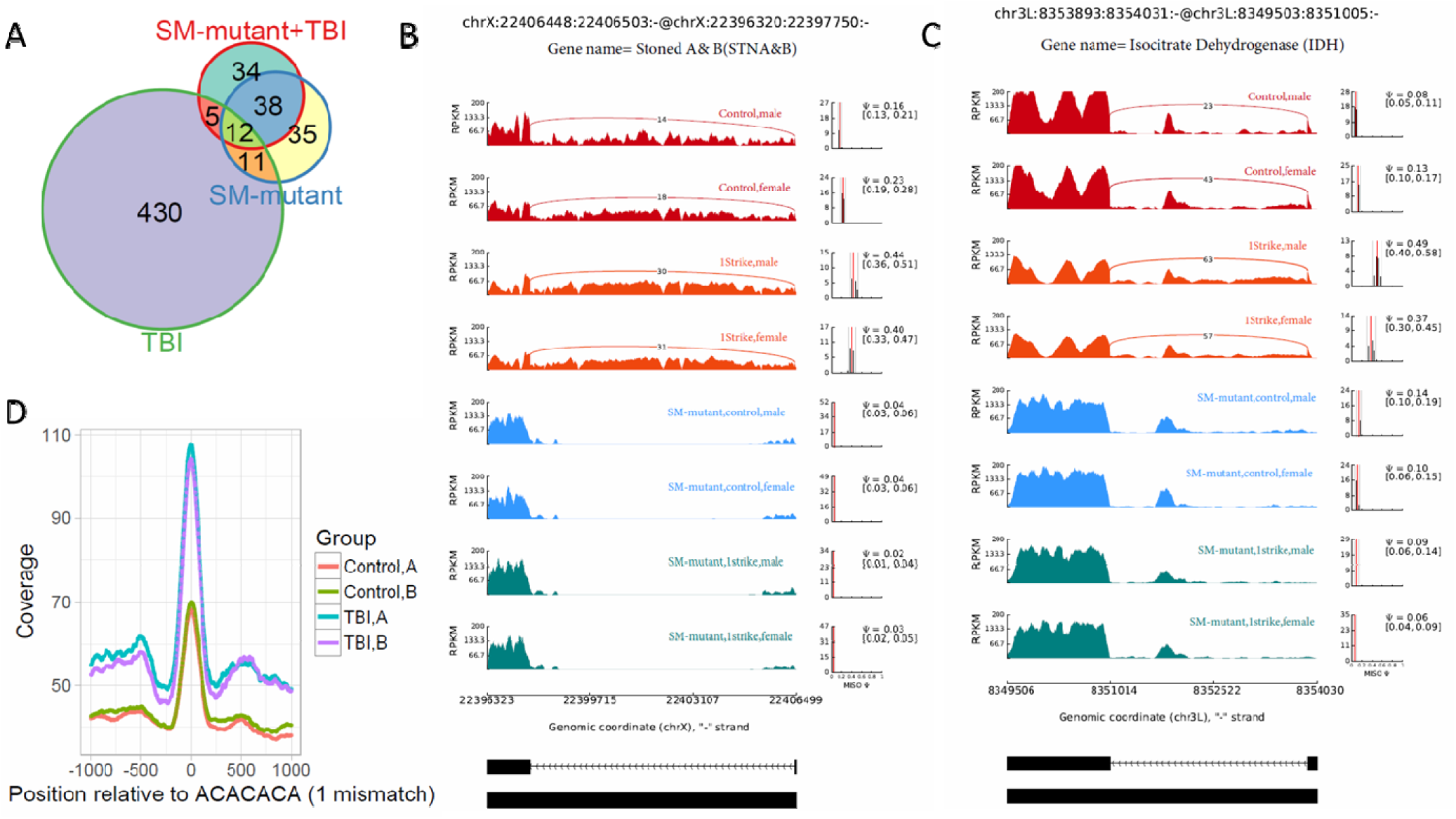
SM is required for intron retention and associated with H3K36me3; A) Overlap between differential RI events detected in TBI, SM-mutant and SM-mutant+TBI samples. B) RI events detected in TBI samples revert back to the non-TBI w1118 controls in SM-mutant and SM-mutant+TBI samples for Stoned A and B. C) RI events detected in TBI samples revert back to the non-TBI w1118 controls in SM-mutant and SM-mutant+TBI samples for Isocitrate Dehydrogenase. D) Enrichment of intronic H3K36me3 peaks centered (±1000) around CA-rich motifs for detected using HOMER (and overlapped with MEDIPS peaks) demonstrate higher normalized coverage for intronic peaks around CA-rich motifs for TBI samples compared to w1118 controls

Additionally, we performed RNA-seq in the sm^4^/Df mutant heads without the TBI and found that the sex-independent differential RI events detected in sm4/Df (-TBI) mutants overlapped with the non-TBI RI events detected in sm^4^/Df +TBI heads (Fig 7A). Examples of this observation are illustrated for STNA and for IDH in Figure 7B and 7C. In STNA the RI events observed in control (w^1118^) flies post-trauma are completely reversed in SM-mutants irrespective of the TBI (Fig 7B). In IDH the RI event is restored to the control (w^1118^) levels in SM-mutant and is non-responsive to TBI (Fig. 7C). Results from this study suggest that SM binding to CA-rich intronic suppressor sites (ISS) may regulate RIs.

### 9. H3K36me3 pattern around SM-binding intron suppressor sites (ISS)

To understand the relationship between SM and H3K36me3 we centered our H3K36me3 intronic peaks on the CA-rich motif (ACACACA) (see Methods). The ACACACA motif positions (100bps centered regions) surrounded by significant H3K36me3 peaks (±1000 bps) for control and 24 hours post-TBI heads are indicated in Figure 7D. We observed that the heads collected 24 hours post-TBI showed higher levels of H3K36me3 in the RIs compared to the controls. Furthermore, ACACACA motif position have notably higher H3K36me3 compared to surrounding regions in both Control and 24 hours post-TBI samples. This observation suggested a possible interaction between H3K36me3 and SM-binding and is discussed further in later sections.

### 10. Modelling

To further understand the association between H3K36me3 and SM-binding in context of RIs we tried to model all intron retention (∆PSI>0) and exclusion events (∆PSI<0) in our TBI samples as a linear function of log fold-change in RPKM of H3K36me3 peaks in TBI compared to control samples, GC frequency, intron length and frequency of ACACACA motif (1 mismatch) near the 3’SS or 5’SS (see Methods). At a p-value ≤ 0.001 we found that the interaction between the intron length, log fold-change in H3K36me3 peaks and frequency of ACACACA motif near 3’SS was predictive of intron retention (∆PSI>0) and showed synergistic affects (Suppl. table 4, Fig 9A). Interestingly, this association was not observed near the 5’SS (Suppl. Table 5, Fig 9B).

### 11. Intron retention in mouse model of spinal cord injury and traumatic brain injury

We have demonstrated that the brain transcriptome undergoes significant changes in the steady-state alternative splicing profile 24 hours after TBI in Drosophila. To determine if the RI plays an important role in regulation of transcriptomic profile in mammalian systems, we performed meta-analysis of 2 published datasets.

In the first study, the authors performed contusive spinal cord injury in 24 female C57BI/6J mice and collected the RNA from spinal tissue 2 days (Acute) and 7 days (Sub-acute) post-SCI. The RNA was sequenced using 100bps paired-end reads (Chen et al. 2013). This mouse SCI model was very similar to our experimental design. First we compared the changes in the exon usage profile 2 days post-SCI and 7 days post-SCI. We observed a significant increase in “exon usage” at 7 days (N= 1136) compared to 2 days (N=456) at a FDR≤0.1 and log2FC≥±2 fold (Fig. 8A and B). This observation was similar to the increase in exon usage we noted 24 hours after-TBI in Drosophila. This suggests the changes in alternative splicing profile in the sub-acute phase of trauma might be conserved across species.

**Figure 8:**
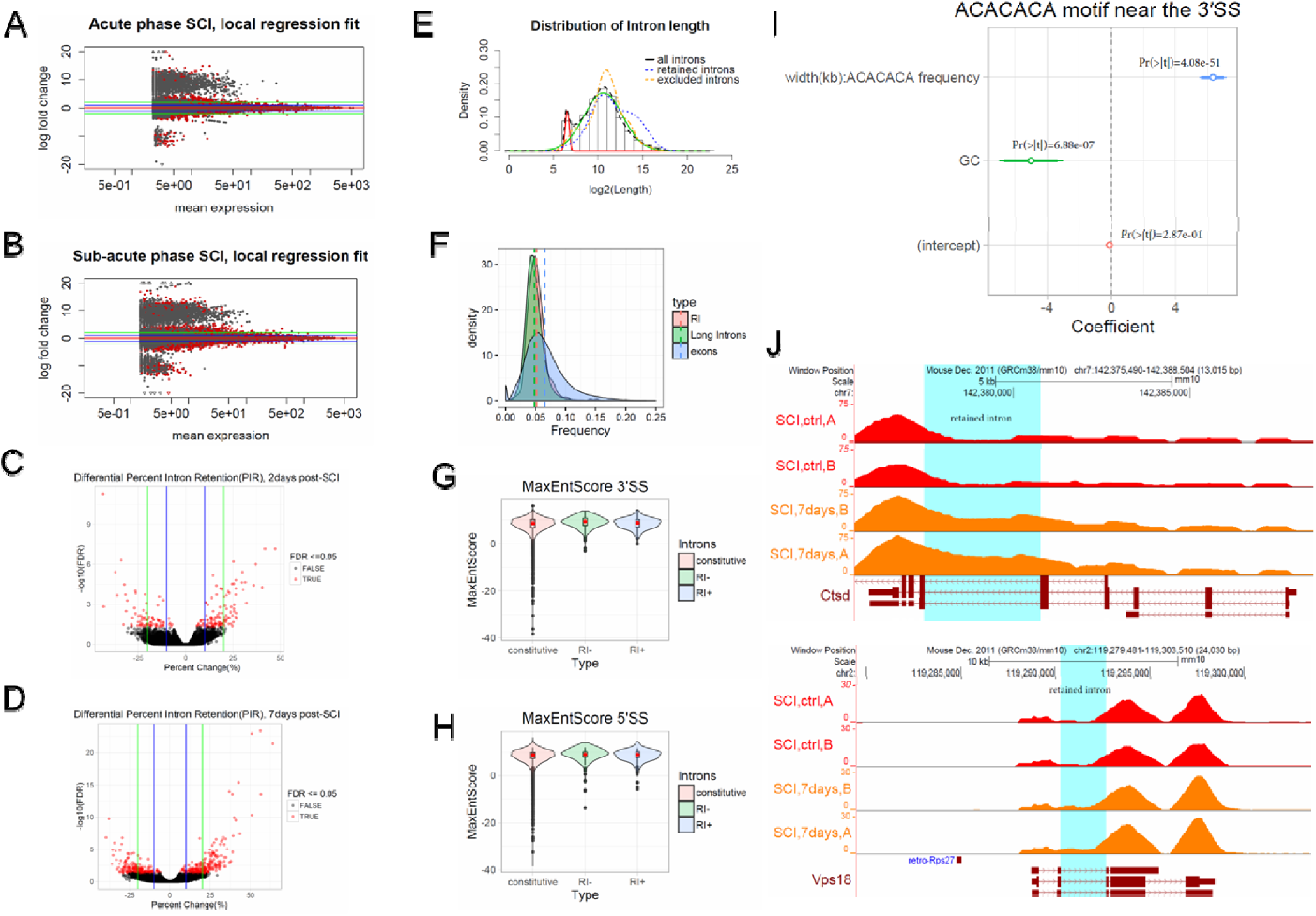
Characterization of alternative splicing events in Mouse model of SCI (Meta-analysis): MA plots differentially used exons for TBI heads collected A) 2days post-SCI (Acute Phase) and B) 7days post-SCI (Sub-Acute phase). Red dots indicate the genes associated with FDR corrected p-value ≤ 0.1 or 10%. Green horizontal line indicate a logFC ≥ ±2 and blue horizontal line indicate a logFC ≥ ±1.C) % change in Percent Intron Retention(PIR) was plotted against –log10 (FDR) for 2days post-SCI. Red dots indicate introns showing significant change in PIR at a FDR corrected p-value ≤ 0.05. Green horizontal line indicate a % change ≥ ±20 and blue horizontal line indicate a % change≥ ±10. D) % change in Percent Intron Retention(PIR) was plotted against –log10 (FDR) for 7days post-SCI. Red dots indicate introns showing significant change in PIR at a FDR corrected p-value ≤ 0.05. Green horizontal line indicate a % change ≥ ±20 and blue horizontal line indicate a % change ≥ ±10. E) Graphical representation of Gaussian mixture model to determine the natural cut-off for calling long introns. Black dotted line represents the log2(intron length) for all introns in the genome. Blue dotted line represents the intron length of retained introns detected for heads collected 7days post-SCI for males and females. Orange dotted line represents the intron length excluded introns detected for heads collected 7days post-SCI for males and females. The red and green line represents the normal distributions fitted using Gaussian mixture model. F) Distribution of GC-content for retained introns + excluded introns (RI), long introns and exons. Dotted lines are the median GC-content. Distribution of Maximum entropy scores (MaxEntScores) for G) 3’SS and H) 5’SS for constitutive introns (+), retained introns (+) (RI+), and retained introns (−) (RI-). I) ∆PIR was modelled as a linear function of GC content + width(Kb): Frequency ACACACA motif(allowing 1 mismatch). “:” indicates an interaction. Coefficients of the model along with respective p-values are illustrated in the figure. All ∆PIR measurements was used irrespective of the FDR corrected p-value. J) Normalized coverage for exons and retained introns for across control and SCI (7days post-SCI) samples for Cathepsin D (CTSD)(panel 1) and vacuole protein sorting 18 (VPS18)(panel 2). Highlighted regions (Cyan) are the retained introns.

RI is widely regarded to be the least prevalent form of AS in higher mammals (Galante et al. 2004; Sakabe and de Souza 2007). However, contrary to the consensus in the field, Braunschweig et al, 2014, reported widespread RI in Human samples. In this study, RI was shown to be involved in the removal of incorrectly spliced transcripts either by nuclear retention or by Nonsense Mediated Decay (NMD) (Braunschweig et al. 2014). For the purpose of detection of mammalian RI, we could not effectively use MISO as the MISO supplied annotation included only 7000 unique introns for the mm10 build of the mice genome. Consequently, analysis for differential splicing events using MISO reported only 2 significant differential RI events 7days post-SCI (Suppl. Fig 7A). Therefore, we adapted the novel approach used by Braunschweig et al, 2014 for detecting differential RI events (Braunschweig et al. 2014). Briefly, we used *spliceSites* package in R/Bioconductor to determine alignment gaps or gap sites in control and post-SCI RNA sequencing. These gap-sites are representative of exon-exon junctions. For quantification of the number of reads aligned to the gap sites, we calculated the Reads Per Million Gapped (RPGM) for all detected gap-sites per sample (see Methods). Then we selected the gap sites for which the RPGM were well correlated (95% CI) between biological replicates and filtered them by RPGM ≥ 5. The resulting output consisted of a list of putative introns (pre-annotated + novel introns) and the number of reads mapping to each respective gap-sites. These reads were classified as “Intron Exclusion Reads”. Among the putative introns, only pre-annotated introns (ENSEMBL introns for mm10) were considered for analysis. The final list included more than 50,000 introns. For these introns, we counted the number of reads that fell within the introns and spanning the intron-exon boundary. These we defined as “Intron Inclusion reads”. Then we calculated Percent RI (PIR) as a ratio of inclusion reads and summation of inclusion and exclusion reads. Finally, to test for significant differences between conditions we performed a *Fisher Exact test*. Further illustration and explanation on PIR calculations are found in the Methods section and Supplemental Figure 7B.

Using PIR we tested RI and exclusion events 2days and 7days post-SCI. After filtering the events by FDR corrected p-value cut-off of 0.05 and ∆PIR ≥ ±10%, we observed 189 events at 2days and 413 events at 7days (Fig. 8C and D). For the RI introns detected 7days post-SCI, we looked at the defining characteristics of the introns such as intron length, GC-content and splice site strength. Using the same strategy used for the Drosophila model of TBI, we calculated the natural cut-off for long introns to be 128 bps (not shown). The majority of the RI/excluded introns detected in the mouse SCI model that were shown to be differentially spliced 7days post-SCI were >128bps in length (Fig 8E). This indicates a conserved retention of long introns in the sub-acute phase of injury. The average GC content of spliced introns was not different from the average GC-content of all long introns in the mouse genome and slightly lower than that of exons (Fig 8F). Similarly, the splice site strength, represented as maximum entropy scores (MaxEntScore) for 5’SS and 3’SS for spliced introns, was not statistically different from constitutive introns (Fig 8G and H). These observations indicate similar characteristics of RI 7days post-SCI in mouse and 24 hours post-TBI in Drosophila. However, only a limited number of genes showing RI showed significant changes in transcript abundance (at an FDR≤0.05 and ∆TPM ≥ ±20) (data not shown).

The 413 RI events detected 7days post-SCI mapped to 375 genes. GO analysis using 6378 genes tested for differential RI as background (see Methods) did not show any specific enrichment of GO classes. Therefore, we performed GO analysis using all ENSEMBL annotated genes for mm10 genomic build as the background gene set. At an FDR ≤ 0.1 we found enrichment of GO categories such as GO:0048471~perinuclear region of cytoplasm, and GO:0000166~nucleotide binding and GO:0005524~ATP binding (Suppl. Fig 2D). This included lysosomal genes implicated in neurodegeneration such as Cathepsin D (CTSD) (Fig 8J top panel) (Cataldo et al. 1995; Koike et al. 2000), and Vacuole protein sorting 18 (VPS18) (Peng et al. 2012a; Peng et al. 2012b) (Fig 8J bottom panel).

Searching for splicing consensus motifs within 300 bps on the intronic side of the 3’SS of RI events, we found the CA-rich regions similar to that detected in the Drosophila TBI experiments (Suppl. Table 6 and 7). Modelling the ∆PIR as a linear function of GC content + width (Kb) to the frequency of the ACACACA motif (allowing 1 mismatch), we demonstrated that a high frequency of ACACACA motif near 3’SS of long intron is associated with RI (see Methods) (Fig 8I). This suggests that the functional association between hnRNPL binding and proper intron processing might be conserved across species. However, further work needs to be done to validate these observations in the mouse models.

The second dataset consisted of sham and TBI cortical and hippocampal samples collected from male C57BL/6J mice. The RNA isolated from these samples was sequenced using 50 bps paired-end sequencing. Filtering the gap-sites per groups by RPGM ≥5 resulted in detection of 2500 novel and pre-annotated introns. The smaller number of gap-site detections in these samples compared with the SCI samples might be attributed to smaller read lengths. Read lengths of 50bps has been shown to be much less efficient in detection of splice junctions compared to 100bps (Chhangawala et al. 2015). In downstream analysis of the cortical samples, we could detect only 2 introns with significant differences (at FDR rate cut-off of 5%) in ∆PIR between sham control and TBI samples. Hierarchical clustering of scaled ∆PIR values showed no segregation of treated and control groups (Suppl. Fig 8A). We observed similar results with the hippocampal transcriptome (Suppl. Fig 8B), further arguing that 50bps read-lengths are insufficient for detection of splice-junctions.

### Discussion

In this study, we explore the alternative splicing changes that take place in the sub-acute phase of mild TBI and their role in regulating survival in Drosophila. The Drosophila TBI model was developed by the Wasserman lab, and was shown to simulate the defining characteristics typical to CHI in humans, such as temporary incapacitation, ataxia, and neurodegeneration (Katzenberger et al. 2013). We adapted this model and modified it to partially ameliorate the impact of the trauma by reducing the force of the impact (Fig 1A). We sequenced RNA from the heads collected 4 hours and 24 hours post-TBI and observed that transcriptomic changes were mainly observed 24 hours after TBI. The transcriptomics changes were driven by alterations in the RNA splicing profile, predominantly intron retention (RI). Therefore, the primary objective for this study was to characterize the RIs and to begin to investigate possible mechanisms regulating these events.

### TBI causes a reduction in mitochondrial metabolism

We observed RIs and consequent reduction in transcript abundance of genes associated with mitochondrial metabolism such as Isocitrate dehydrogenase (IDH), Enolase (ENO), Pyruvate Kinase (PYK) and Aconitase (ACON). Retention of long introns in these transcripts can introduce pre-mature termination codons (PTMs) and target the RNA for degradation by Nonsense Mediated Decay (NMD) (Wong et al. 2013; Braunschweig et al. 2014). These genes with TBI-induced RIs encode for critical regulators in the Krebs/TCA cycle. For examples, ACON catalyzes the conversion of cis-aconitate to D-isocitrate (Beinert et al. 1996) and IDH catalyzes the conversion of D-isocitrate to α-Ketoglurate (Gabriel et al. 1986; Yen and Schenkein 2012).

To characterize the impact of TBI on mitochondrial metabolism, we measured the cytochrome *c* oxidase (COX) activity in control and 24 hours post-TBI heads. COX is the last enzyme of the respiratory electron transport chain and is responsible for conversion of molecular oxygen to water. During this process, the proton pumps transport protons across the inner mitochondrial membrane, generating an electrochemical gradient, which is utilized by ATP synthase to produce ATP. Under normal conditions, COX is downregulated by allosteric inhibition by ATP (Arnold and Kadenbach 1997). Under oxidative stress conditions, the allosteric inhibition is nullified, resulting in hyperpolarization of the membrane potential and production of ROS. We observed an increased COX activity 24 hours post-TBI, which suggests a loss of ATP-dependent inhibition and increase in mitochondrial stress (Kadenbach et al. 2004). Consistent with the COX activity measurements, we also observed a decrease in ATP availability. Repair and tissue remodeling post-TBI is an energy expensive process. Coupled with the dysregulation of Krebs/TCA cycle genes, TBI may further lead to a reduction of ATP levels and loss of allosteric inhibition of COX. This result suggests a regulatory link between RI in TCA cycle genes post-TBI and a consequent decrease in mitochondrial metabolism.

Downregulation of mitochondrial metabolism is an adaptive and neuro-protective mechanism in neurodegenerative disorders. For example, restoration of mitochondrial function by sustained exposure to creatinine (5 mM creatinine for 7 days) (Andres et al. 2005) resulted in an increase in ATP levels and oxidation/reduction activity but a decrease in cell viability of Down syndrome (DS)-derived astrocytic cells (Helguera et al. 2013). This suggested that an increase in mitochondrial activity in neurodegenerative conditions might paradoxically have harmful rather than beneficial effects.

We believe that the genes associated with the TCA cycle in our experiments that show an increase in RI 24 hours after TBI, such as IDH and ACON, could be “critical epicenters” that regulate neuroprotection in neurodegenerative conditions. Dysregulation of steady state RNA levels of these genes can significantly impact the production of a critical metabolite that is involved in epigenetic regulation, such as α-Ketoglutarate(α-KG) (Chia et al. 2011). α-KG is an important co-factor in regulating all dioxygenase reactions in the cell including H3K36me3 demethylation by the Jumonji-family of histone demethylases (Klose and Zhang 2007). In a recent study by Carry et al, 2015, direct perturbation of α-KG/succinate levels was sufficient to cause changes in H3K27me3 and DNA methylation in mouse embryonic stem cells, suggesting α-KG is an important metabolic effector of epigenetic modification by both histone and DNA demethylases (Carey et al. 2015). Based on the above arguments, we believe that RI and the consequent decrease in transcript abundance of IDH and ACON provide suggestive evidence of the regulatory link between mitochondrial metabolism and epigenetic processes. We speculate that RI post-TBI result in decreases levels of α-KG, the direct product of the IDH enzyme, thereby leading to decreased activity of KDM4A and accumulation of H3K36me3 in intronic regions. Increased H3K36me3 in coordination with splicing factors such as SM (hnRNPL) can cause even more increased levels of RI and further downregulation of mitochondrial metabolism to facilitate recovery of damaged neurons post-trauma (Fig 9D).

**Figure 9:**
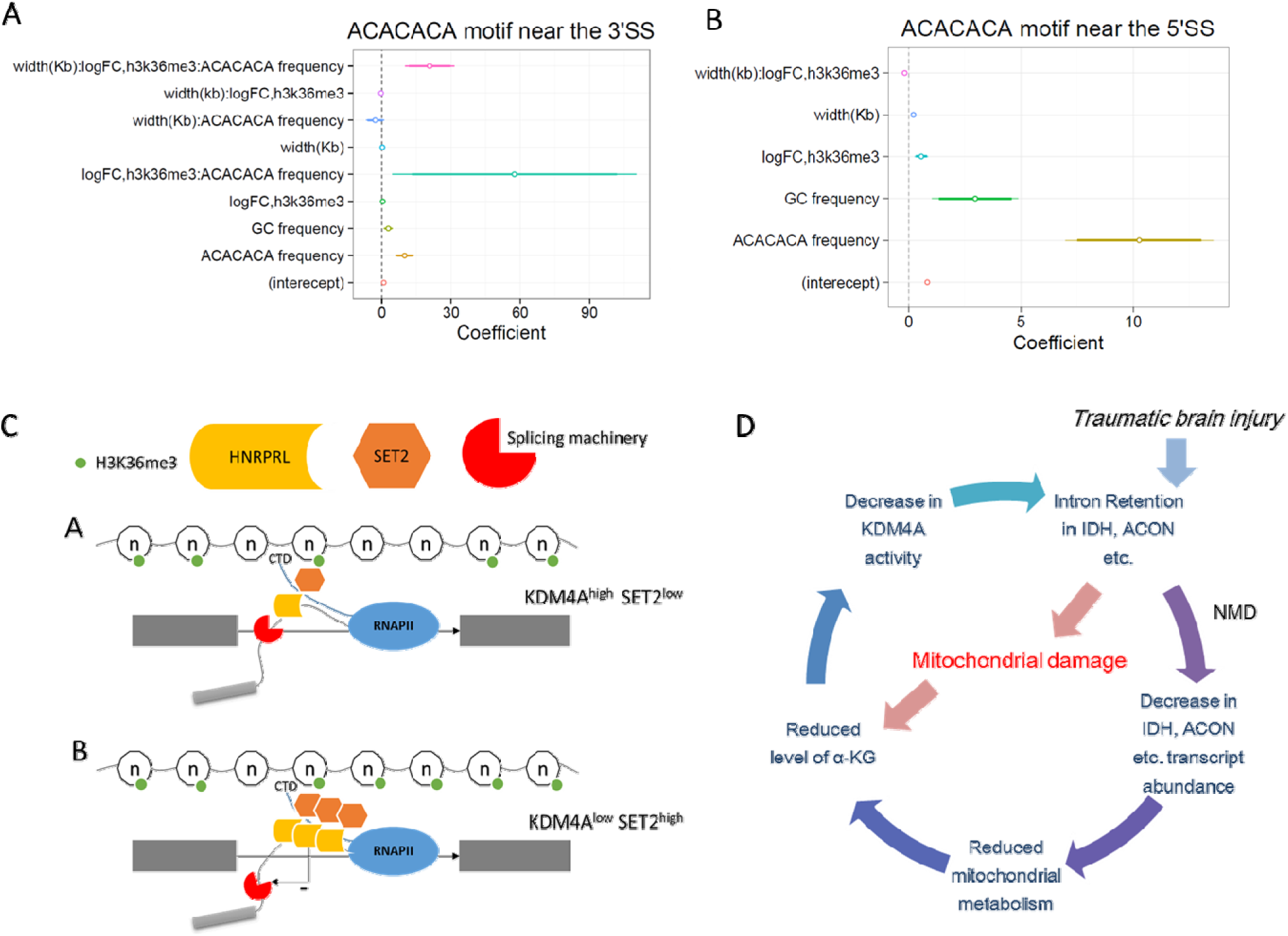
Model for regulation of intron retention in Drosophila in the sub-acute phase of traumatic brain injury. A) Contribution of factors in regulation of intron retention. GC frequency= GC frequency, width(Kb)= intron length (Kb), logFC= RPKM(TBI)/RPKM(Control) and ACACACA frequency= Frequency of ACACACA motif near the 3’SS allowing for 1 mismatch. Width(Kb):logFC:ACACACA frequency show synergistic effect on intron retention (RI). B) Contribution of factors in regulation of intron retention. GC frequency= GC frequency, width(Kb)= intron length (Kb), logFC= RPKM(TBI)/RPKM(Control) and ACACACA frequency= Frequency of ACACACA motif near the 5’SS allowing for 1 mismatch. C) Top panel: Under normal conditions the introns are marked by lower levels of H3K36me3 compared to the flanking exons. This allows the elongating polymerase and the associated splicing machinery to recognize it as an intron. The binding of hnRNP-L/Smooth to intronic splicing suppressor sites under normal conditions is limited and cause only a basal level of intron retention. Bottom panel: Post-TBI the level of H3K36me3 in introns increases and now it resembles an exon therefore is spliced in by the splicing machinery. This is a direct result in recruitment of hnRNP-L/Smooth and SET2 complex to intronic splicing suppressor sites. D) Hypothesized mechanism of regulation of mitochondrial metabolism through intron retention in genes involved in TCA cycle and Glycolysis.

### The RNA binding protein SM interferes with splicing of long introns after TBI

Based on the motif search within the annotated RIs post-TBI we found a specific enrichment of CA-rich sequences near the 5’SS and 3’SS. The CA-rich motif has been shown to bind to the RNA binding protein Smooth (SM), which shares approx. 42% identity with the hnRNPL class of splicing regulators (Suppl. Fig 10) (Ray et al. 2013). The hnRNPL splicing factors are versatile in their function and can regulate a wide variety of splicing events, including skipping of cassette type exons and suppression of variable exons. Notably, hnRNPL knockdown and hnRNPL/hnRNPLL double knockdown causes RI in CD55 and STRA6 genes in HeLa cell lines, suggesting that they may be required for efficient processing of introns (Hung et al. 2008). The Drosophila genome encodes a homologue of the hnRNPL protein, known as Smooth (SM). SM contains 3 protein motifs corresponding to RRM (RNA binding) domains which share approximately 42%, 25% and 42% identity with RRM domains of human hnRNPL proteins (Suppl. Figure 10).

To determine the relationship between SM protein and intron retention in our TBI model, we investigated *sm*-mutant (*sm4*/Df) hemizygous flies (Karpen and Spradling 1992; Layalle et al. 2005; Draper et al. 2009). We used the *sm4*/Df mutant instead of traditional RNAi or CRISPR-KO lines because *sm*-mutant flies have well characterized, reproducible and easily observable phenotypes. For example, the *sm*-mutants have an average lifespan of 30 days, compared with 60-90 days for wild type flies, due to defective feeding behavior (Layalle et al. 2005; Draper et al. 2009). This feeding behavior phenotype is attributed to the lack of chemosensory axon arborization in the leg neuromeres (Layalle et al. 2005; Draper et al. 2009). Behavioral analyses of *sm*-mutant flies also indicated progressive muscle degeneration leading to pronounced motor impairments, which may also contribute to a decreased life span for these flies (Draper et al. 2009). Subjecting the *sm4*/Df mutants to TBI resulted in the loss of RIs observed 24 hours’ post-trauma in control w^1118^ flies. In some cases, for example in the STNA/B 1^st^ intron, the basal level of RI that is observed in the absence of TBI was also eliminated. This result provides strong evidence that SM (hnRNPL) binding to intronic splicing suppressor (ISS) sites is necessary for RI 24 hours post-TBI.

Our result that SM-binding to introns causes RI is not consistent with the observation reported by Hung and colleagues in HeLa cells, who showed that SM is required for proper processing of CD55 and STRA6 introns (Hung et al. 2008). However, it is worth noting that splicing factors such as SM have multiple binding sites along the genes – some in introns and some in exons. The effects of SM on RNA splicing may vary depending upon their binding to intronic or exonic sites. For instance, in our TBI model, we hypothesize that SM (hnRNPL) is most likely being recruited to intronic sites after TBI, and that this binding results in increased levels of RI. An increase in SM-binding to RI introns can be investigated by using RNA-immunoprecipitation (RIP)-sequencing using antibodies to SM. However, the lack of ChIP-grade antibodies for Drosophila SM, is a major limitation. We acknowledge this as one of the caveats of this study and we are currently attempting to address this issue by analyzing viable GFP-tagged SM mutant lines. As the *sm4*/Df flies are short lived, SM-mediated RI might be protective in nature. Neuroprotection by SM-mediated RI is most likely occur by temporarily decreasing the translation of mitochondrial enzymes and, thereby decreasing mitochondrial metabolic output.

### The histone modification H3K36me3 interferes with splicing of long introns after TBI

In previous sections, using direct calculation of Splicing Rates (SR) from RNA-Seq data, we have argued that a majority of splicing in single transcript genes in Drosophila takes place co-transcriptionally. This is in agreement with previous observations. For example, S2 cells expressing a slow polymerase mutant show a greater efficiency of splicing; *i.e*., a more efficient processing of introns (Khodor et al. 2011). This demonstrates that the speed of the elongating polymerase has a profound impact on proper intron processing. This also suggests that alternative splicing in Drosophila takes place co-transcriptionally. If splicing is coupled to transcription, then local histone modifications might be involved in the regulation of splicing.

One of the modifications that has been implicated in alternative splicing regulation is Histone 3 lysine 36 tri-methylation (H3K36me3). Yoh et al, 2008 reported that SETD2 histone methyl-transferase (HMT) is recruited to the C-terminal domain of the elongating RNA polymerase by SPT6, an elongation factor. This facilitates the addition of H3K36me3 onto the genome co-transcriptionally (Yoh et al. 2008). Furthermore, H3K36me3 marks the gene bodies of actively transcribing genes and is depleted from the introns and enriched in the exons in C.elegans and mouse (Kolasinska-Zwierz et al. 2009). H3K36me3 is also depleted in alternative exons compared to flanking constitutive exons (Kolasinska-Zwierz et al. 2009). Interestingly, we found that depletion of H3K36me3 from alternate exons compared to flanking constitutive exons is also observed in Drosophila, suggesting that association between H3K36me3 and alternative splicing might be an evolutionarily conserved (Suppl Fig 6 and Fig 6 A-D).

Direct measurement of H3K36me3 within RIs using ChIP-Seq demonstrated that a subset of RIs showed an increase in H3K36me3 modification in 24 hours post-TBI heads. This result was further substantiated by our KDM4A-RNAi studies in 3^rd^ instar larvae. KDM4A is the predominant demethylase of H3K36me3 in Drosophila. Therefore, if H3K36me3 is associated with RIs, then using KDM4A-RNAi in 3rd instar larvae, we should be able to partially mimic some of the RIs found in 24 hours post-TBI heads. In agreement with this hypothesis, we observed some of the major RIs found 24 hours post-TBI in KDM4A-RNAi 3rd instar larvae, including RIs in two of the genes associated with mitochondrial metabolism (IDH and PYK) and neuronal transport protein STNA and STNB.

H3K36me3 has been reported to indirectly interact with splicing factors such as PTB and SRSF1 and, thereby, cause exon skipping in the fibroblast growth factor receptor (FGFR2) gene (Luco et al. 2010; Pradeepa et al. 2012; Zhou et al. 2014). A study by Yuan et al, 2009 reported hnRNPL to be an integral part of the lysine trimethylase 3A (KTM3A) complex (Yuan et al. 2009). KMT3A (also known as HYPB or hSET2) is a histone methyltransferase specifically shown to increase H3K36me3 levels in mammals and Drosophila (Bell et al. 2007; Edmunds et al. 2008). The authors demonstrated that RNA interference against KMT3A or hnRNPL down-regulates exclusively the H3K36me3 mark in HeLa cells. This suggests that hnRNPL might be required for hSET2 methyltransferase activity. Interestingly remapping H3K36me3 peaks around ACACACA motif showed high enrichment of H3K36me3 peaks at the motif-position and lower enrichment in the surrounding 1000 bps regions in both control and 24 hours post-TBI samples (Suppl. Fig 5C). High enrichment of H3K36me3 in intronic regions around SM-binding motifs (ACACACA) is indicative of localized H3K36 methyltransferase activity in this region. This observation within limitation does support the interaction between SM (hnRNPL) and SET2 HMT, therefore we believe that this possible association merit future experimental validation.

Mathematical modelling also revealed the interaction between frequency of ACACACA motif near 3’SS, width of the intron and logFC in H3K36me3 between control and 24 hours post-TBI heads within introns is a positive predictor of intron retention (RIs) (Figure 9A and 9B). Therefore, our study so far provides strong suggestive evidence indicating involvement of SM (hnRNPL) and H3K36me3 in regulation of intron processing in TBI.

Based on evidence from previous studies combined with our study, we hypothesize that, 24 hours’ post-trauma, there is an increase in recruitment of SM (hnRNPL) binding proteins to the 3’SS of long introns. This leads to recruitment of SET2 HMT to the ACACACA sites and increases the levels of H3K36me3 within these introns (Fig 9C). Simultaneously, an increase in mitochondrial stress post-TBI reduces the availability of α-KG, and, thereby, causes a decrease in KDM4A histone demethylase activity (Fig 9D). Therefore, the disruption of balance between the SETD2 histone methyltransferase and KDM4A histone demethylase activities contribute to an increase in H3K36me3 within long introns (Fig 9C and 9D). As exons are known to be marked by higher levels of H3K36me3 compared to introns (Kolasinska-Zwierz et al. 2009) (Fig 6B, C, and D), we speculate that this increase in intronic H3K36me3 levels post-trauma by the SM, SET2 and KDM4A might be responsible for misrecognition of introns as exons by the splicing machinery and consequent intron retention.

The Drosophila model of TBI poses several interesting and as-of-yet unresolved questions. For example, (1) what provides the initial signal for SM (hnRNPL) to bind to long introns post-trauma; (2) does SM (hnRNPL) physically associate with the SET2 HMT and contribute to increase H3K36me3 within long introns post-trauma; (3) is α-KG the rate limiting metabolite that controls H3K36me3 demethylation through modulation of KDM4A activity; and (4) how is the dynamic balance between H3K36 methylation and demethylation regulated? In future studies, we will continue to explore these questions and further our understanding of regulation of alternative splicing in sub-acute phase of traumatic brain injury.

### Intron retention of long introns also occurs in mouse models of neural trauma

Functional conservation especially in response to neuronal damage in Drosophila and mammalian systems is well studied. For example, over-expression of human Tau proteins in fly heads causes progressive neurodegeneration and behavioral, motor and olfactory learning deficits (Beharry et al. 2013). Also, abnormal CAG repeat expansion of ATXN3 gene causes neurodegeneration in mammalian and fly models, and are ubiquitinated in mammalian cells and in Drosophila (Tsou et al. 2013). Furthermore, neuronal support systems reported to play an important role in recovery from neuronal damage in mammalian systems are also present in Drosophila. For example, the Drosophila brain contains so-called ensheathing and reticular glial cells that demonstrate ameboid morphology and have been reported to be involved in the clearance of cellular debris by endocytosis (MacDonald et al. 2006; Doherty et al. 2009). Therefore, in essence, ensheathing and reticular glial cells function as microglia homologues in Drosophila.

Though parallels exist at the behavioral and cellular level, studies have demonstrated a strong conservation in the regulation of splicing events and exon–intron architecture between higher mammals and Drosophila. For example, both Drosophila and mammalian alternatively spliced exons are often flanked by long introns, and the length of the upstream intron seems to play a more important role in exon selection than downstream introns (Fox-Walsh et al. 2005). This association has been reported to be much more evident in mammals, which suggests a positive selection for specific exon–intron architectures throughout evolution (Fox-Walsh et al. 2005).

Our meta-analysis of a mouse model of Spinal Cord Injury (SCI) showed a similar increase in RI in the sub-acute phase of TBI/7days post-SCI and a specific enrichment of the CA-rich motif around 3’SS of the RI introns, similar to what we observe in Drosophila. It is important to note that the mouse dataset consisted of RNA isolated from peripheral nervous tissue, whereas, our data was from fly heads. Additionally, SCI is caused in their studies by direct physical trauma to an exposed spinal cord, and is therefore probably more severe compared to our Drosophila model of TBI. Even considering the significant differences between our study and the mouse SCI studies, the presence of hnRNPL binding sites near the 3’SS of RIs post-SCI is intriguing and suggests an evolutionarily conserved mechanism of intron retention post-neuronal damage. Further investigation using both Drosophila and mouse models of TBI are required to determine the extent of extent of evolutionary conservation and the precise mechanism of protection against these injuries.

## Material and Methods

### 1. Sample for survival estimation and exploratory RNA sequencing

Sample consisted of 0 to 5days old w^1118^ flies. These flies were collected in sturdy plastic vials and subjected to Traumatic brain injury using the High Impact trauma (HIT) device. For the survival assay, the male and female flies were separated and 25 flies per vial were used for each condition (control, 1 strike, 2 strike). The flies were subjected to TBI, at a spring deflection of approx. 45° to ameliorate the impact of trauma. The 2^nd^ strike was performed after a recovery time of 5mins. These flies were maintained till ~98% of the flies in the control vial were deceased. Survival estimations and standard error calculation was done using R/Bioconductor. For RNA sequencing analysis heads from 50 males and 50 females flies for control, 1 strike and 2 strikes were pulled off manually one at a time and immediately transferred to *RNAlater (500µl)*. Fly head collection was done at 2 time-points 4 hours and 24 hours. The male and female were used as biological replicates. More explanation of study design for RNA-sequencing studies is provided in later sections.

### 2. Bulk preparation of fly heads

0-5days old w^1118^ flies were transferred to 50 ml tubes on ice. Approximately 3 ml of flies was collected per tube. These flies were snap frozen by dipping them in liquid nitrogen and vortexed vigorously to detach the head (for 3mins; 15 secs per turn). The vial was dipped in liquid nitrogen between consecutive turns to prevent them from thawing. The vortexed flies were passed through a sieve 720µm pore size. This allowed the separation of the bodies from the detached heads. The detached heads were collected on another sieve 410µm pore size. This sieve allowed the separation of heads from other detached body parts. The fly heads were transferred to 2ml Eppendorf tubes and stored in −80°C till further use.

### 3. Sample for Sub-cellular fractionation

0-5days old w^1118^ flies were collected in bulk and separated into 2 batches. Batch 1 was used as control and prepared for subcellular fractionation. The fly heads were collected in *RNAlater (500µl)* to prevent RNA degradation. Batch 2 was transferred to plastic vial and subjected to 1 strike and collected after 24 hours in *RNA-later* (500µl). Samples were stored in −20°C prior to fractionation. Approximately, 500µg of heads were collected per condition in 2ml Eppendorf tubes.

### 4. Sample for ChIP-sequencing

0-5days old w^1118^ flies were collected in bulk and separated into 2 batches. Batch 1 was used as control and prepared for tissue homogenization and DNA isolation. The fly heads were collected in 1XPBS (phosphate buffer saline) containing protease halt cocktail (10µl per ml of PBS). Batch 2 was transferred to plastic vial and subjected to 1 strike and collected after 24 hours in 1XPBS (+ protease halt cocktail). Approximately, 500µg of heads were collected per condition in 2ml Eppendorf tubes.

### 5. 5. Sample for RNA-sequencing 3^rd^ instar larvae and SM-mutant flies

Most of the gene knockdowns were performed by crossing da-Gal4 flies, which have ubiquitous expression, to UAS-RNAi flies using the following stocks from the Bloomington, Indiana stock center: y[1] sc[*] v[1]; Py[+t7.7] v[+t1.8]=TRiP.HMS01304 attP2 (expresses dsRNA for RNAi of Kdm4A (FBgn0033233) under UAS control); y[1] v[1]; Py[+t7.7] v[+t1.8]=TRiP.HMS02273 attP40 (expresses dsRNA for RNAi of Idh (FBgn0001248) under UAS control); P{GAL4-da.G32), which expresses Gal4 in all cells; y[1] v[1]; Py[+t7.7] v[+t1.8]=TRiP.JF01320 attP2 (Expresses dsRNA for RNAi of Kdm2 (FBgn0037659) under UAS control); y[1] sc[*] v[1]; Py[+t7.7] v[+t1.8]=TRiP.HMS00488 attP2 (Expresses dsRNA for RNAi of CG7200 (FBgn0032671) under UAS control); y[1] sc[*] v[1]; Py[+t7.7] v[+t1.8]=TRiP.HMS00775 attP2 (Expresses dsRNA for RNAi of CG8165 (FBgn0037703) under UAS control); and y[1] sc[*] v[1]; Py[+t7.7] v[+t1.8]=TRiP.HMS00575 attP2 (Expresses dsRNA for RNAi of Utx (FBgn0260749) under UAS control). The smooth-RNAi lines were not viable when crossed to da-Gal4, so instead we used a hypomorphic allele over a deficiency in a hemizygous combination; cn[1] P{PZ} sm[05338]/CyO; ry[506] (aka. sm [4]) and w[1118]; Df(2R)BSC820, P+PBac w[+mC]=XP3.WH3 BSC820/SM6a.

### 6. Tissue homogenization

For tissue homogenization 4 stainless steel 3.2mm beads (Qiagen cat no. 69990) was added to 2ml Eppendorf tubes. The samples were subjected to homogenization using TissuelyserLT (Qiagen) using the following setting; 50 oscillations (1/s) for 2min. The homogenized samples were centrifuged using a low-speed bench top centrifuge and supernatant containing cells were collected for downstream processing.

### 7. Subcellular fractionation

The *RNAlater* has a specific gravity greater than cells therefore it is difficult to recover the complete homogenate for prepared samples. To optimize the recovery, the *RNAlater* was diluted using 1XPBS (1:2). Subcellular fractionation was carried out using RNA Subcellular Isolation Kit (Active motif; 25501). The nuclear fraction was further treated with DNase to degrade contaminating DNA using DNase treatment kit (Active motif, 25503). The isolated RNA was measured using Trinean DropSense96 Spectrophotometer.

### 8. RNA isolation

RNA was isolated as described in the attached Qiagen EZ1 RNA Handbook, 3rd ed. 06/2012, page 34, for "Any Type of Tissue" using the EZ1 RNA Universal Tissue Kit with TissueLyser II homogenization/disruption and optional DNase digestion. RNA was quantified by UV Spec on the Trinean DropSense96 Spectrophotometer. RNA quality assessment were performed using the Agilent RNA ScreenTape on the Agilent 2200 TapeStation.

### 9. Chromatin immunoprecipitation

The homogenized tissue collected was fixed by incubating with 16% Formaldehyde (final concentration 1%) for 10 min at RT. Then 10XGlycine (final concentration 1X) was added and samples were incubated at RT for 5 mins, to quench the activity of Formaldehyde. After incubation the samples were centrifuged at 3000g for 5min and supernatant was removed to obtain cell pellets. The formaldehyde fixed cell pellets were washed 2 times with PBS (1X)+ HC (3000g for 5min) and then reconstituted in Membrane extraction buffer + HC (Pierce Chromatin Prep module, 26158). Cell pellets were broken up by pipetting with a p200 and vortexing for 15 secs and incubated on ice for 15 min. The lysed cells were recovered by centrifugation (9000g for 3min at 4°C) and reconstituted in 200 or 300µl of Nuclease free water (depending upon the size of the pellet, decided arbitrarily). Sonication was carried out using Covaris S2 Series 2 Focused Ultra-sonicator at intensity=5, duty cycle: 10%, 200cycle/burst, for 5-8 mins (depending upon size of the pellet and presence of debris). The sonicated nuclei were recovered using centrifugation (10000g for 5min) and reconstituted in Nuclear extraction buffer + HC (Pierce Chromatin Prep module, 26158). The nuclei were incubated on ice for 15 min which 15sec of vortexing every 5 min. The nuclei were centrifuged at 9000g for 5 min and supernatant containing cleaved chromatin was collected for downstream analyses. The DNA concentrations were measured using Qbit. Chromatin bound DNA was collected from mass preps (N=4) of control and TBI heads (24 hours post-TBI) and pooled and redistributed for immunoprecipitation using Anti-Histone H3 (tri methyl K36) antibody - ChIP Grade (ab9050). We used total of 0.5µg/reaction of DNA for control and TBI heads. The efficiency of the sonication and quality of IP was assessed using 2200 TapeStation System. The IP’ed DNA was sequenced using 100bps paired-end reads in replicates. Quality control metric from sequencing runs were estimated using HOMER (http://homer.salk.edu/homer).

### 10. Mitochondrial profiling

A. **Sample:** 0-5days old w^1118^ flies were transferred to 50 ml tubes on ice. Approximately 3 ml of flies was collected per tube. These flies were snap frozen by dipping them in liquid nitrogen and vortexed vigorously to detach the head (for 3mins; 15 sec per turn). The vial was dipped in liquid nitrogen between consecutive turns to prevent them from thawing. The vortex flies were passed through a sieve 720µm pore size. This allowed the separation of the bodies from the detached heads. The detached heads were collected on another sieve 410µm pore size. This sieve allowed the separation of heads from other detached body parts. The fly heads were transferred to 2ml Eppendorf tubes kept on dry ice. Then they the stored at −80°C until measurement. We had a total of 16 control samples (0.038 ± 0.013 gm/tube) and 15 24-hours post-TBI samples (0.018 ± 0.004 gm/tube).
B. **Cytochrome *c* oxidase (COX) activity measurements**: To test the effect of TBI on electron transport chain function, COX activity measurements were performed as described Lee et al, 2005 and 2009 (Lee et al. 2005; Lee et al. 2009). Briefly, fly heads were collected and stored frozen at −80 ºC until measurements were performed. COX activity was measured with a micro Clark-type oxygen electrode in a closed chamber (Oxygraph system, Hansatech) at 25°C. Frozen samples were minced in 10 mM K-HEPES, 40 mM KCL, 1% Tween-20, 2 mM EGTA, 1 mM Na-vanadate, 1 mM PMSF, 1 µM oligomycin) and solubilized by sonication for 5 seconds for 2 times. The supernatant was collected, and COX activity measured in the presence 20 cow heart cytochrome c (Sigma) and 20 mM ascorbic acid as reductant. Oxygen consumption was recorded on a computer and analyzed with the Oxygraph software. Protein concentration was determined with the DC protein assay kit (Bio-Rad, Hercules, CA, USA). COX specific activity was defined as consumed O_2_ (nmol)/min/mg total protein and reported as % of control.
C. **Measurement of total ATP concentration:** ATP levels were determined via the bioluminescent Method (HS II kit, Roche Applied Science) in conjunction with the boiling Method as described (Samavati et al. 2008).

### 11. RNA sequencing and ChIP sequencing

Library for RNA sequencing was prepared as described in the TruSeq Sranded Total RNA Sample Preparation Guide, Rev. E, Oct. 2013, using the TruSeq Stranded Total RNA LT (w/ RiboZero Human/Mouse/Rat) -Set A library kit. 100 bps paired-end libraries for RNA and ChIP sequencing were sequencing on the Illumina HiSeq 2500 following cBot clustering.

### 12. Statistical analysis

#### A. RNA sequencing dataset used for meta-analysis

For the first study i.e. RNA sequencing of spinal cord tissue from mouse subjected to spinal contusion 2days or 7days post-injury, the data was directly downloaded from Gene Expression Omnibus (GEO accession number: GSE45376). For the second study i.e. RNA sequencing of hippocampal and cortical tissue from mice subjected to mTBI, the raw .fastq files were obtained from the authors.

For the first mouse dataset the authors performed contusive spinal cord injury in 24 female C57BI/6J mice and collected the RNA from spinal tissue 2days (Acute) and 7days (Sub-acute) post-SCI. The RNA was sequenced using 100bps paired end reads. This model was very similar to our experimental design. The second dataset consisted of sham and TBI cortical and hippocampal samples collected from Male C57BL/6J mice in biological triplicates. The RNA isolated from these samples was sequenced using 50 bps paired-end sequencing.

#### B. RNA-sequencing alignments

Prior to alignment the sequencing adaptors were trimmed using FASTQ Toolkit v1.0 in Illumina Basespace. FastQC was used to generate quality metrics for assessment of .fastq files. RNA sequencing alignment was done using TopHat v2.1.0 or HISAT 0.1.6-beta using *default* alignment parameters. After alignment the mapped reads was further filtered by mappability score (MAPQ≥ 10). The quality controlled .bam files were sorted by genomic position of the reads using *samtools-0.1.19*. PCR duplicate reads were processed and removed using *rmdup* function (options -S) in *samtools*. The sorted duplicate removed .bam files were further assessed and visualized using Integrative Genome Viewer (https://www.broadinstitute.org/igv/v1.4). The Mouse and Drosophila dataset were both processed using this pipeline. For Drosophila Melanogaster UCSC genomic build *dm3* was used for read alignments. For Mus Musculus UCSC genomic build *mm10* was used for read alignment.

#### C. ChIP-Sequencing alignments

All ChIP sequencing alignments were done using *Bowtie2* using the dm3 build of the Drosophila genome as reference. After alignment the mapped reads was further filtered by mappability score (MAPQ≥ 10). The quality controlled. bam files were sorted by genomic position of the reads using *samtools*. PCR duplicate reads were processed and removed using rmdup function (options -S) in *samtools*. The sorted duplicate removed .bam files were further assessed and visualized using Integrative Genome Viewer (https://www.broadinstitute.org/igv/v1.4). The tag density and Quality metrics such as GC content and genomic nucleotide frequencies relative (-50, 50) to the 5’ end of ChIP-fragment were calculated using HOMER (http://homer.salk.edu/homer). (Suppl. figure 8)

#### D. Gene expression analysis

Reads from the processed .bam files were overlapped with ENSEMBL exon annotation extracted from UCSC in R/Bioconductor (Packages; *GenomicAlignments, GenomicFeatures*). The dm3 genomic-build was used for Drosophila datasets and mm10 genomic-build was used for the Mouse dataset. The read count per exon was computed using the preset “Union” mode. (Refer to *summarizeoverlaps*). The exon counts were next combined to give the gene read counts. Differential gene expression analysis from computed read counts was carried out using DESEQ2 in R/Bioconductor. The read count and DESEQ2 parameters used were identical between the Drosophila and mouse dataset.

For Drosophila model of TBI RNA were prepared in bulk for 50 males and 50 females for 3 separate conditions; control, 1 strike and 2 strikes at 2 time points; 4 hours and 24 hours. This meant that for the exploratory analysis we had a total of 12 samples. The negative binomial distribution (see DESEQ2 vignette) (Love et al. 2014) was fitted using treatment conditions; control, 1 strike and 2 strikes as classifiers and sex of the flies and time of collection as categorical covariates (read distribution ~ conditions + sex + time of collection). This enabled us to control for any potential sources of gene expression variations which may arise due to the sex of the flies. Additionally, this allowed us to use the male and the female w^1118^ flies were used as biological replicates. Male and female w^1118^ non-TBI flies collected at 4 hours and 24 hours were used as controls for all differential expression estimations. The differentially expressed genes were further filtered using FDR and log2FC cut-off which are further discussed in the results section.

#### E. Exon usage analysis

Number of aligned reads was counted within disjoint exonic bins rather than exons using the “Union” mode of read-counting in *summarizeoverlaps*. Then DEXSEQ was used to determine relative exon-usage while controlling for overall gene expression. Relative exon usage can be defined as follows;

Exon usage= # transcripts from the gene that contain target exon/# all transcripts from the gene

The read counting and DEXSEQ parameters used were identical between the Drosophila and mouse dataset.

For Drosophila TBI the negative binomial distribution for modelling read counts (see DEXSEQ vignette) was fitted using treatment conditions; control, 1 strike and 2 strikes as classifiers and sex of the flies and time of collection as categorical covariates. For each target gene DEXSEQ models read distribution as a function of ~conditions + exon + sex:exon+ time:exon + conditions:exon, where “:” indicate existence of possible interaction/correlations between covariates. Then this complete model is compared against a *null* model; ~conditions + exon + sex:exon+ time:exon, to determine the effect of treatment conditions of exon usage. For the mouse SCI dataset DEXSEQ analysis was performed using ~ conditions + exon + conditions:exon as the complete model and ~conditions + exons as the *partial/null* model. Further details on DEXSEQ can be found in the package vignette and the paper(Anders et al. 2012).

#### F. Mixture of Isoforms (MISO) analysis

Exon usage exclusively relies on reads mapping to exons. However, much more accurate estimation of alternative splicing changes can be inferred from intronic and junction reads. For this purpose, we used Mixture of isoform (MISO) to estimate the Percent Spliced in (PSI) value for each annotated splicing feature. Annotations for splicing features were provided by modENCODE consortium and classified into 5 representative classes; Alternative 3’SS, Alternative 5’SS, Skipped exons, mutually exclusive exons, and Retained introns. The difference in PSI value (∆PSI) between control and treatment (TBI) samples were estimated using Bayesian factor analysis. The comparisons were further filtered using a ∆PSI cut-off of 0.05 or 5%, Bayesian factor ≥ 10 and number of exclusion and inclusion reads ≥ 10 (*see MISO documentation*). The significant events common between males and females were selected and correlated to give the final list of sex-independent splicing changes. As majority of the significant events were retained introns downstream annotation and visualization was done only with retained intron genes. The retained introns were overlapped with maximum overhang length of 10 bps with introns annotation obtained from the ENSEMBL genes UCSC dm3 build of the genome and annotated with respective gene and transcript annotation. Visualization of MISO results was done using 2 independent approaches. In the first approach, the log_10_(RPKM) (reads per kilobase per million) for respective splicing features were plotted using sashimi plot (https://miso.readthedocs.io/en/fastmiso/). In the second approach, the normalized coverage was for exonic and intronic regions was calculated using HOMER (http://homer.salk.edu/homer).

#### G. Characterization of introns

Introns and their respective lengths in base-pairs were obtained for ENSEMBL genes from UCSC dm3 build of the Drosophila genome. The lengths were log transformed (log_2_) and their density distribution was determined. Then the density distribution was modelled as a mixture of N=2 normal distribution using *Gaussian mixture model*. This allowed us to determine the natural cut-off for long introns. The distributions inferred from the model were plotted using R/Bioconductor. The GC content was determining directly from the .fasta sequence corresponding to the dm3 build of the Drosophila genome and plotted using *ggplot2* in R/Bioconductor. Maximum entropy scores (MaxEntScores), was used to discriminate between weak and strong splice sites flanking the retained introns. Briefly, short sequence motif around 5’SS and 3’SS for retained introns was collected depending upon the strand for the gene. MaxEntScores was calculated using the *score5* and *score3* functions in R/Bioconductor (Package *spliceSites*) and was confirmed using the online version (http://genes.mit.edu/burgelab/maxent/Xmaxentscan_scoreseq.html). Intron characterization for retained introns in mouse dataset was carried out using exactly the same pipeline. We used mouse ENSEMBL genome UCSC build mm10 for the mouse datasets.

#### H. Calculating Transcripts per million (TPM)

For determining relationship between transcript abundance and RI we calculated the Transcript Per Million (TPM) for all transcripts containing retained introns for all samples.

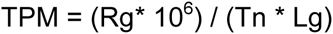

Where Rg is total read counts for the transcript, Tn is the sum of all length normalized total transcript read counts, and length is the length of the transcript in bps. Following TPM calculations, we tested from significant differences in TPM between representative sample groups using *Fisher Exact Test* (Package; *edgeR*) and filtered the transcripts by FDR corrected p-value of 0.05 and ∆TPM ≥ ±20. The ∆TPM were plotted against –log_10_(FDR) in R/Bioconductor. The same Method for TPM calculation was used for Drosophila and mouse datasets.

#### I. Calculating Splicing rate (SR)

Co-transcriptional splicing rate or Splicing rate (SR) was calculated following the same principle proposed by Wickramasinghe et al, 2015(Wickramasinghe et al. 2015). The splicing rate is defined as the normalized read counts of the first intron versus the last intron of each transcript. Calculations in equation form are as follows;

f_i_= C_i_/L_i_ where C_i_ is the read count for last introns and L_i_ is the length of last intron

f_k_= C_k_/L_k_ where C_k_ is the read count for first introns and L_k_ is the length of first intron

f_i-1_= C_i-1_/L_i-1_ where C_i-1_ is the read count for last but one exon and L_i-1_ is the length of last but one exon

f_k+1_= C_k+1_/L_k+1_ where C_k+1_ is the read count for 2^nd^ exon and L_k+1_ is the length of 2^nd^ exon

F_i_= f_i_/(f_i-1_X N) where N= library size of the sample

F_k_= f_k_/(f_k+1_X N) where N= library size of the sample

SR= F_k/_F_i_ where SR= splicing rate

Transcripts selected for SR calculation was selected based on following criteria; belongs to a single transcript gene, number of exon for the transcript/gene ≥ 4.

#### J. Meta-analysis of ChIP sequencing datasets

*Exon-trio analysis* (ETA) for ChIP sequencing data was adapted from Kolasinska-Zwierz, et al, 2009(Kolasinska-Zwierz et al. 2009). Briefly, we used ENSEMBL gene model annotation from BioMart to create random sets of Constitutive and Alternative exon trios. The constitutive trio (CE trios) consisted of 3 adjacent CE. The alternative trio (AE trios) consisted of 1 AE flanked by 2 CE in a constitutive-alternative-constitutive configuration. We selected an average of unique 1-3 trios per candidate gene. Then we counted the number of reads mapping to exon belonging to pre-defined trios using *MEDIPS* package from R/Bioconductor. We compared each ChIP set to their inputs within 200bps sliding windows and selected exons which showed at least +2fold enrichment of ChIP peaks over input at an FDR corrected p-value ≤ 0.05. For the list of enriched exons, we filtered them such that all 3 members of the trio are present and significantly enriched. We labelled these member as 1= exon1, 2=exon2, 3=exon3 based on their genomic organization and position. Finally, we compared the tag counts (read counts) for each member of the constitutive or alternative trio to each other using ANOVA followed by Tukey-HSD test in R/Bioconductor.

The relative enrichment of H3K36me3 peaks in exons and introns where done using the *MEDIPS* package from R/Bioconductor. Briefly, the differential tag densities within 200 bp regions were estimated using MEDIPS. Regions which showed a ≥ + 2-fold enrichment at an FDR corrected p-value ≤ 0.05 over input, was considered as significant peaks. The target peaks were overlapped with regions of interest (ROIs). Criteria for selecting ROIs are as follows; introns and exons containing significant peaks were ≥200 bps in length, all introns and exons of the transcripts had significant peaks in them (N=3752). Reads Per Kilobase per Million (RPKM) for respective ROIs were plotted with *ggplo2* in R/Bioconductor.

#### K. Analysis of ChIP-sequencing data

We used 0.5µg DNA per ChIP reaction. This is a relatively small amount compared to recommended 50 to 100µg. Therefore, we expected relatively less enrichment of H3K36me3 peaks with large background signal. We tried multiple peak calling strategies including the more commonly used peak caller such as MACS1.4.2. We found that combination of MEDIPS in R/Bioconductor and HOMER was more suited for low enrichment high background data. We sequenced our sample using 100bps paired-end reads for more accurate estimation of fragment length and used replicates to control for biases.

We estimated 2 important quality metrics GC% bias and Genomic Nucleotide Frequency relative to read positions using HOMER. These are illustrated in Supplemental Figure 9. These quality metrics were well within expected values. Following the quality control for our H3K36me3 ChIP-seq datasets we determined significant peaks within 200bps regions using MEDIPS. We implemented 2 additional controls in peak calls; firstly, replaced all reads which map to exactly the same start and end positions by only one representative read. Secondly, we specified that the ChIP-seq data used for peak calling was paired end, for more accurate estimation of fragment length. Significant peak calls were made against input controls for TBI and Control samples and filtered by an FDR corrected P-value ≤ 0.1 or 10%. Additionally, significant peaks were called for H3K36me3 modENCODE data using an FDR corrected P-value ≤ 0.05 and log2FC ≥ +2fold. The significant peaks called for w^1118^ control, TBI, and H3K36me3 modENCODE samples where aggregated and averaged across all differential RIs detected in TBI samples. The RIs with significant peaks was overlapped between the w^1118^ control, TBI, and H3K36me3 modENCODE samples using *Vennerable* in R/Bioconductor. For the common RIs the RPKM (Reads per Kilo-base per Million) were compared between w^1118^ control and TBI using Welch 2sample t-test.

For determining the H3K36me3 levels around the CA-rich motifs we reanalyzed our H3K36me3 data for w^1118^ control and TBI using HOMER. We called peaks against their respective input control using the following parameters; <u>“*-fdr 0.1 -P 0.1 -F 0 -L 0 -LP 0.1 -style factor”.*</u> The peaks detected were overlapped with MEDIPS peaks detected with intronic region for w^1118^ control and TBI (only peaks with minimum overlap 10 bases were kept). We re-centered these peaks around CA-rich motifs (ACACACA allowing 1 mismatch) and estimated the read coverage ±1000 bps around the CA-rich regions. The results were plotted using *ggplot2* in R/Bioconductor.

#### L. Splicing factor discovery

We limited our motif discovery to intronic splicing sites i.e. motifs within RIs. The median intron size of retained introns was ~3000bps (min=115, max= 36560). Therefore, only retained introns with size ≥600 bps (421/458) was considered for motif analysis. For potential splicing factor binding motifs, 300bp regions were collected from the intronic side of the 5’SS and 3’SS. Sequences for these target regions were obtained and Motif enrichment analysis was carried out using DREME (Discriminative Regular Expression Motif Elicitation). DREME uses reshuffled input sequences as control to calculate motif enrichment. There are wide varieties of motif enrichment analysis software available publicly, which use varied mathematical models for enrichment test. Therefore, we wanted to confirm our finding in DREME using different analysis strategy; MotifRG (R/Bioconductor). We searched for enrichment of motifs against the randomly selected control sets of ~300bps windows from the dm3 build of the Drosophila genome. The enrichment analysis was run with 5 bootstrapping test to estimate score variance and filtered by bootstrap P-value ≤ 0.1 (MotifRG default parameters). Only the Top 10 results from MotifRG were considered.

#### M. Calculating RI in mammalian datasets

For mouse SCI and mTBI, the Percent RI (PIR) calculations were adapted from Braunschweig et al, 2015.

PIR= (Number of inclusion reads X 100)/ (Number of exclusion reads + Number of inclusion reads)

For calculating the *number of inclusion reads*, the first step was to define a list of target regions for read counting. We selected all annotated introns per transcripts from ENSMEBL (UCSC mm10 build) as our target regions. For this pre-defined set of targets, we counted the reads using *summarizeoverlaps* function in “Union” mode. *Inclusion reads* is comprised of 2 types of reads; mid-intron reads i.e. reads which lie within the intron and exon-intron junction reads which span the edges of the intron. “Union” mode of read counting accounts for the position of the read with respect to the target regions and any overlapping feature. Therefore, it is efficiently able to assign mid-intron and exon-intron junction reads to the respective targets. A better illustration of “Union” mode of counting reads can be found in Suppl. Fig 7A.

Exclusion reads on the other hand are equivalent to junction reads i.e. reads which maps across exon-exon junctions. Using the utilities provided in the spliceSites package in R/Bioconductor we mapped the gap-sites corresponding to the junction reads for each sample. These gap-sites are representative of introns. For each gap-sites aka introns we counted the total number of junction read alignment and calculate RPGM values.

RPGM= (Number of aligns per splice site X 10^6)/Number of gapped aligns per probe.

Then we filtered our gap-sites such that they have an RPGM ≥ 5 and are well correlated (~95% CI) across replicated RNA-sequencing samples. The final list consisted of a set of novel and pre-annotated introns/gap-sites. We further filtered our curated intron sets and only considered gap-sites (introns) which overlapped with the targets used for counting the *inclusion reads* (maximum overhang length ≤ 3 bp), i.e. pre-annotated introns. The number of reads corresponding to these final set of gap sites are defined as *exclusion reads*.

Finally, for each intron/sample we calculated the PIR values as described before, and compared the PIR values between respective groups of samples using *Fisher Exact Test.* The results were further filtered using ∆PIR ≥ ±10% and FDR corrected p-value≤ 0.05. For visualization of intron exclusion/retention the Normalized coverage was estimated using HOMER and plotted using Genome Browser.

#### N. Modelling of intron retention in Drosophila and SCI datasets

For modelling intron retention, we selected all the intron retention and exclusion events irrespective of the ∆PSI and Bayes factor values and converted them in categorical variables (∆PSI > 0 ~ TRUE, ∆PSI < 0 ~ FALSE). For determining the frequency of ACACACA motif we divided the introns in equal halves and counted the number of ACACACA motif allowing 1 mismatch at any base position. The frequency was calculated as follows.

ACACACA frequency= ACACACA count *2/intron length

Then we modelled categorical ∆PSI as a function of GC frequency, intron length (width), Log-fold change in mean RPKM of H3K36me3 peaks mapping to RI in 24 hours post-TBI samples compared to controls and frequency of ACACACA motif (1 mismatch) near the 3’SS or 5’SS.

Cat(∆PSI) ~ GC frequency + width(Kb)*log2FC*ACACACA frequency

To determine the best model, we used *step* function to select a formula-based model by AIC. Following the selection of the best model the coefficients were extracted and plots were made using ggplot2.

For modelling intron retention and exclusion in mouse SCI datasets we selected all RI/exclusion events (N=30460) irrespective of their FDR corrected p-value. For these events we divided the introns in equal halves and counted the number of ACACACA motif allowing 1 mismatch at any base position. Only sequences near 3’SS was chosen for further study as motif analysis placed the ACACACA motif near the 3’SS in mouse model of SCI. The frequency was calculated as follows.

ACACACA frequency= ACACACA count *2/intron length

Then we modelled ∆PIR (continuous) as a linear function of GC frequency, intron length (width) and frequency of ACACACA motif (1 mismatch) near the 3’SS. We selected the best model based on AIC using the *step* function in R/Bioconductor. The best model was as follows:

∆PIR~ GC frequency + width(Kb)* ACACACA frequency

#### O. Gene ontology analysis

Gene ontology (GO) analysis was performed using DAVID bioinformatics resources (https://david-d.ncifcrf.gov). The background gene set for GO analysis were selected as follows;

1. gene expression; 5923 genes which were expressed in both controls and 24 hours post-TBI samples were used as background,
2. exon-usage; 9056 genes containing well represented exons (counts/exon ≥ 2) was used as background,
3. RI, Drosophila, dm3; 4209 genes containing RI events detected both in males and females were used as controls
4. PIR 7 days post-SCI, mm10; 6378 genes which were used for differential PIR test were used as background.

If no, GO categories were found to be enriched than the all ENSEMBL genes (dm3 or mm10) were used as background. The Fold enrichment (over background) was plotted against the -log_10_(FDR) for visualization of significant GO categories in ggplot2. The GO categories with FDR ≤ 0.1 was labelled in red.

#### P. Data repository

All sequencing data has been submitted to Gene Expression Omnibus (GSE…….)

## Acknowledgements

The research is funded to D.M.R. by NIH grants R01 ES012933 and P30 ES020957.

## References

Acosta SA, Tajiri N, de la Pena I, Bastawrous M, Sanberg PR, Kaneko Y, Borlongan CV. 2015. Alpha-synuclein as a pathological link between chronic traumatic brain injury and Parkinson’s disease. Journal of cellular physiology 230(5): 1024–1032.

Anders S, Reyes A, Huber W. 2012. Detecting differential usage of exons from RNA-seq data. Genome research 22(10): 2008–2017.

Andres RH, Ducray AD, Huber AW, Perez-Bouza A, Krebs SH, Schlattner U, Seiler RW, Wallimann T, Widmer HR. 2005. Effects of creatine treatment on survival and differentiation of GABA-ergic neurons in cultured striatal tissue. Journal of neurochemistry 95(1): 33–45.

Arnold S, Kadenbach B. 1997. Cell respiration is controlled by ATP, an allosteric inhibitor of cytochrome-c oxidase. European journal of biochemistry /FEBS 249(1):1 350–354.

Arnold S, Kadenbach B. 1999. The intramitochondrial ATP/ADP-ratio controls cytochrome c oxidase activity allosterically. FEBS letters 443(2): 105–108.

Beharry C, Alaniz ME, Alonso Adel C. 2013. Expression of Alzheimer-like pathological human tau induces a behavioral motor and olfactory learning deficit in Drosophila melanogaster. Journal of Alzheimer’s disease: JAD 37(3): 539–550.

Beinert H, Kennedy MC, Stout CD. 1996. Aconitase as Ironminus signSulfur Protein, Enzyme, and Iron-Regulatory Protein. Chemical reviews 96(7): 2335–2374.

Bell O, Wirbelauer C, Hild M, Scharf AN, Schwaiger M, MacAlpine DM, Zilbermann F, van Leeuwen F, Bell SP, Imhof A et al. 2007. Localized H3K36 methylation states define histone H4K16 acetylation during transcriptional elongation in Drosophila. The EMBO journal 26(24): 4974–4984.

Braunschweig U, Barbosa-Morais NL, Pan Q, Nachman EN, Alipanahi B, Gonatopoulos-Pournatzis T, Frey B, Irimia M, Blencowe BJ. 2014. Widespread intron retention in mammals functionally tunes transcriptomes. Genome research 24(11): 1774–1786.

Broglio SP, Eckner JT, Paulson HL, Kutcher JS. 2012. Cognitive decline and aging: the role of concussive and subconcussive impacts. Exercise and sport sciences reviews 40(3): 138–144.

Browne KD, Chen XH, Meaney DF, Smith DH. 2011. Mild traumatic brain injury and diffuse axonal injury in swine. Journal of neurotrauma 28(9): 1747–1755.

Carey BW, Finley LW, Cross JR, Allis CD, Thompson CB. 2015. Intracellular alpha-ketoglutarate maintains the pluripotency of embryonic stem cells. Nature 518(7539): 413–416.

Cataldo AM, Barnett JL, Berman SA, Li J, Quarless S, Bursztajn S, Lippa C, Nixon RA. 1995. Gene expression and cellular content of cathepsin D in Alzheimer’s disease brain: evidence for early up-regulation of the endosomal-lysosomal system. Neuron 14(3): 671–680.

Chang YF, Imam JS, Wilkinson MF. 2007. The nonsense-mediated decay RNA surveillance pathway. Annual review of biochemistry 76: 51–74.

Chen K, Deng S, Lu H, Zheng Y, Yang G, Kim D, Cao Q, Wu JQ. 2013. RNA-seq characterization of spinal cord injury transcriptome in acute/subacute phases: a resource for understanding the pathology at the systems level. PloS one 8(8): e72567.

Chhangawala S, Rudy G, Mason CE, Rosenfeld JA. 2015. The impact of read length on quantification of differentially expressed genes and splice junction detection. Genome biology 16: 131.

Chia N, Wang L, Lu X, Senut MC, Brenner C, Ruden DM. 2011. Hypothesis: environmental regulation of 5-hydroxymethylcytosine by oxidative stress. Epigenetics: official journal of the DNA Methylation Society 6(7): 853–856.

Copur O, Muller J. 2013. The histone H3-K27 demethylase Utx regulates HOX gene expression in Drosophila in a temporally restricted manner. Development 140(16): 3478–3485.

Crona F, Dahlberg O, Lundberg LE, Larsson J, Mannervik M. 2013. Gene regulation by the lysine demethylase KDM4A in Drosophila. Developmental biology 373(2): 453–463.

Daneshvar DH, Goldstein LE, Kiernan PT, Stein TD, McKee AC. 2015. Post-traumatic neurodegeneration and chronic traumatic encephalopathy. Molecular and cellular neurosciences 66(Pt B): 81–90.

Dardiotis E, Fountas KN, Dardioti M, Xiromerisiou G, Kapsalaki E, Tasiou A, Hadjigeorgiou GM. 2010. Genetic association studies in patients with traumatic brain injury. Neurosurgical focus 28(1): E9.

Doherty J, Logan MA, Tasdemir OE, Freeman MR. 2009. Ensheathing glia function as phagocytes in the adult Drosophila brain. The Journal of neuroscience: the official journal of the Society for Neuroscience 29(15): 4768–4781.

Draper I, Tabaka ME, Jackson FR, Salomon RN, Kopin AS. 2009. The evolutionarily conserved RNA binding protein SMOOTH is essential for maintaining normal muscle function. Fly 3(4): 235–246.

Edmunds JW, Mahadevan LC, Clayton AL. 2008. Dynamic histone H3 methylation during gene induction: HYPB/Setd2 mediates all H3K36 trimethylation. The EMBO journal 27(2): 406–420.

Fox-Walsh KL, Dou Y, Lam BJ, Hung SP, Baldi PF, Hertel KJ. 2005. The architecture of pre-mRNAs affects mechanisms of splice-site pairing. Proceedings of the National Academy of Sciences of the United States of America 102(45): 16176–16181.

Gabriel JL, Zervos PR, Plaut GW. 1986. Activity of purified NAD-specific isocitrate dehydrogenase at modulator and substrate concentrations approximating conditions in mitochondria. Metabolism: clinical and experimental 35(7): 661–667.

Galante PA, Sakabe NJ, Kirschbaum-Slager N, de Souza SJ. 2004. Detection and evaluation of intron retention events in the human transcriptome. Rna 10(5): 757–765.

Gennarelli TA, Thibault LE, Adams JH, Graham DI, Thompson CJ, Marcincin RP. 1982. Diffuse axonal injury and traumatic coma in the primate. Annals of neurology 12(6): 564–574.

Giza CC, Hovda DA. 2014. The new neurometabolic cascade of concussion. Neurosurgery 75 Suppl 4: S24–33.

Hall RC, Hall RC, Chapman MJ. 2005. Definition, diagnosis, and forensic implications of postconcussional syndrome. Psychosomatics 46(3): 195–202.

Helguera P, Seiglie J, Rodriguez J, Hanna M, Helguera G, Busciglio J. 2013. Adaptive downregulation of mitochondrial function in down syndrome. Cell metabolism 17(1): 132–140.

Herz HM, Morgan M, Gao X, Jackson J, Rickels R, Swanson SK, Florens L, Washburn MP, Eissenberg JC, Shilatifard A. 2014. Histone H3 lysine-to-methionine mutants as a paradigm to study chromatin signaling. Science 345(6200): 1065–1070.

Huang X, Dixit VM. 2011. Cross talk between ubiquitination and demethylation. Molecular and cellular biology 31(18): 3682–3683.

Hung LH, Heiner M, Hui J, Schreiner S, Benes V, Bindereif A. 2008. Diverse roles of hnRNP L in mammalian mRNA processing: a combined microarray and RNAi analysis. Rna 14(2): 284–296.

Isken O, Maquat LE. 2007. Quality control of eukaryotic mRNA: safeguarding cells from abnormal mRNA function. Genes & development 21(15): 1833–1856.

Jackson TC, Du L, Janesko-Feldman K, Vagni VA, Dezfulian C, Poloyac SM, Jackson EK, Clark RS, Kochanek PM. 2015. The nuclear splicing factor RNA binding motif 5 promotes caspase activation in human neuronal cells, and increases after traumatic brain injury in mice. Journal of cerebral blood flow and metabolism: official journal of the International Society of Cerebral Blood Flow and Metabolism 35(4): 655–666.

Kadenbach B, Arnold S, Lee I, Huttemann M. 2004. The possible role of cytochrome c oxidase in stress-induced apoptosis and degenerative diseases. Biochimica et biophysica acta 1655(1-3): 400–408.

Karpen GH, Spradling AC. 1992. Analysis of subtelomeric heterochromatin in the Drosophila minichromosome Dp1187 by single P element insertional mutagenesis. Genetics 132(3): 737–753.

Katz Y, Wang ET, Airoldi EM, Burge CB. 2010. Analysis and design of RNA sequencing experiments for identifying isoform regulation. Nature methods 7(12): 1009–1015.

Katzenberger RJ, Loewen CA, Wassarman DR, Petersen AJ, Ganetzky B, Wassarman DA. 2013. A Drosophila model of closed head traumatic brain injury. Proceedings of the National Academy of Sciences of the United States of America 110(44): E4152–4159.

Khodor YL, Rodriguez J, Abruzzi KC, Tang CH, Marr MT, 2nd, Rosbash M. 2011. Nascent-seq indicates widespread cotranscriptional pre-mRNA splicing in Drosophila. Genes & development 25(23): 2502–2512.

Klose RJ, Zhang Y. 2007. Regulation of histone methylation by demethylimination and demethylation. Nature reviews Molecular cell biology 8(4): 307–318.

Koike M, Nakanishi H, Saftig P, Ezaki J, Isahara K, Ohsawa Y, Schulz-Schaeffer W, Watanabe T, Waguri S, Kametaka S et al. 2000. Cathepsin D deficiency induces lysosomal storage with ceroid lipofuscin in mouse CNS neurons. The Journal of neuroscience: the official journal of the Society for Neuroscience 20(18): 6898–6906.

Kolasinska-Zwierz P, Down T, Latorre I, Liu T, Liu XS, Ahringer J. 2009. Differential chromatin marking of introns and expressed exons by H3K36me3. Nature genetics 41(3): 376–381.

Kunz WS, Kudin A, Vielhaber S, Elger CE, Attardi G, Villani G. 2000. Flux control of cytochrome c oxidase in human skeletal muscle. The Journal of biological chemistry 275(36): 27741–27745

Layalle S, Coessens E, Ghysen A, Dambly-Chaudiere C. 2005. Smooth, a hnRNP encoding gene, controls axonal navigation in Drosophila. Genes to cells: devoted to molecular & cellular mechanisms 10(2): 119–125.

Lee I, Salomon AR, Ficarro S, Mathes I, Lottspeich F, Grossman LI, Huttemann M. 2005. cAMP-dependent tyrosine phosphorylation of subunit I inhibits cytochrome c oxidase activity. The Journal of biological chemistry 280(7): 6094–6100.

Lee I, Salomon AR, Yu K, Samavati L, Pecina P, Pecinova A, Huttemann M. 2009. Isolation of regulatory-competent, phosphorylated cytochrome C oxidase. Methods in enzymology 457: 193–210.

Li Y, Rao X, Mattox WW, Amos CI, Liu B. 2015. RNA-Seq Analysis of Differential Splice Junction Usage and Intron Retentions by DEXSeq. PloS one 10(9): e0136653.

Lin CH, Paulson A, Abmayr SM, Workman JL. 2012. HP1a targets the Drosophila KDM4A demethylase to a subset of heterochromatic genes to regulate H3K36me3 levels. PloS one 7(6): e39758.

Ling H, Hardy J, Zetterberg H. 2015. Neurological consequences of traumatic brain injuries in sports. Molecular and cellular neurosciences 66(Pt B): 114–122.

Loane DJ, Kumar A, Stoica BA, Cabatbat R, Faden AI. 2014. Progressive neurodegeneration after experimental brain trauma: association with chronic microglial activation. Journal of neuropathology and experimental neurology 73(1): 14–29.

Lorbeck MT, Singh N, Zervos A, Dhatta M, Lapchenko M, Yang C, Elefant F. 2010. The histone demethylase Dmel\Kdm4A controls genes required for life span and male-specific sex determination in Drosophila. Gene 450(1-2): 8–17.

Love MI, Huber W, Anders S. 2014. Moderated estimation of fold change and dispersion for RNA-seq data with DESeq2. Genome biology 15(12): 550.

Luco RF, Pan Q, Tominaga K, Blencowe BJ, Pereira-Smith OM, Misteli T. 2010. Regulation of alternative splicing by histone modifications. Science 327(5968): 996–1000.

MacDonald JM, Beach MG, Porpiglia E, Sheehan AE, Watts RJ, Freeman MR. 2006. The Drosophila cell corpse engulfment receptor Draper mediates glial clearance of severed axons. Neuron 50(6): 869–881.

McKee AC, Robinson ME. 2014. Military-related traumatic brain injury and neurodegeneration. Alzheimer’s & dementia: the journal of the Alzheimer’s Association 10(3 Suppl): S242–253.

Peng C, Yan S, Ye J, Shen L, Xu T, Tao W. 2012a. Vps18 deficiency inhibits dendritogenesis in Purkinje cells by blocking the lysosomal degradation of Lysyl Oxidase. Biochemical and biophysical research communications 423(4): 715–720.

Peng C, Ye J, Yan S, Kong S, Shen Y, Li C, Li Q, Zheng Y, Deng K, Xu T et al. 2012b. Ablation of vacuole protein sorting 18 (Vps18) gene leads to neurodegeneration and impaired neuronal migration by disrupting multiple vesicle transport pathways to lysosomes. The Journal of biological chemistry 287(39): 32861–32873.

Pradeepa MM, Sutherland HG, Ule J, Grimes GR, Bickmore WA. 2012. Psip1/Ledgf p52 binds methylated histone H3K36 and splicing factors and contributes to the regulation of alternative splicing. PLoS genetics 8(5): e1002717.

Ray D, Kazan H, Cook KB, Weirauch MT, Najafabadi HS, Li X, Gueroussov S, Albu M, Zheng H, Yang A et al. 2013. A compendium of RNA-binding motifs for decoding gene regulation. Nature 499(7457): 172–177.

Sakabe NJ, de Souza SJ. 2007. Sequence features responsible for intron retention in human. BMC genomics 8: 59.

Samavati L, Lee I, Mathes I, Lottspeich F, Huttemann M. 2008. Tumor necrosis factor alpha inhibits oxidative phosphorylation through tyrosine phosphorylation at subunit I of cytochrome c oxidase. The Journal of biological chemistry 283(30): 21134–21144.

Song Y, Sretavan D, Salegio EA, Berg J, Huang X, Cheng T, Xiong X, Meltzer S, Han C, Nguyen TT et al. 2015. Regulation of axon regeneration by the RNA repair and splicing pathway. Nature neuroscience 18(6): 817–825.

Sullivan S, Friess SH, Ralston J, Smith C, Propert KJ, Rapp PE, Margulies SS. 2013. Behavioral deficits and axonal injury persistence after rotational head injury are direction dependent. Journal of neurotrauma 30(7): 538–545.

Tsou WL, Burr AA, Ouyang M, Blount JR, Scaglione KM, Todi SV. 2013. Ubiquitination regulates the neuroprotective function of the deubiquitinase ataxin-3 in vivo. The Journal of biological chemistry 288(48): 34460–34469.

Tsurumi A, Dutta P, Shang R, Yan SJ, Li WX. 2013. Drosophila Kdm4 demethylases in histone H3 lysine 9 demethylation and ecdysteroid signaling. Sci Rep 3: 2894.

Unterharnscheidt F. 1995. A neurologist’s reflections on boxing. I: Impact mechanics in boxing and injuries other than central nervous system damage. Revista de neurologia 23(121): 661–674.

Villani G, Greco M, Papa S, Attardi G. 1998. Low reserve of cytochrome c oxidase capacity in vivo in the respiratory chain of a variety of human cell types. The Journal of biological chemistry 273(48): 31829–31836.

Wickramasinghe VO, Gonzalez-Porta M, Perera D, Bartolozzi AR, Sibley CR, Hallegger M, Ule J, Marioni JC, Venkitaraman AR. 2015. Regulation of constitutive and alternative mRNA splicing across the human transcriptome by PRPF8 is determined by 5’ splice site strength. Genome biology 16: 201.

Wong JJ, Ritchie W, Ebner OA, Selbach M, Wong JW, Huang Y, Gao D, Pinello N, Gonzalez M, Baidya K et al. 2013. Orchestrated intron retention regulates normal granulocyte differentiation. Cell 154(3): 583–595.

Yen KE, Schenkein DP. 2012. Cancer-associated isocitrate dehydrogenase mutations. The oncologist 17(1): 5–8.

Yeo G, Burge CB. 2004. Maximum entropy modeling of short sequence motifs with applications to RNA splicing signals. Journal of computational biology: a journal of computational molecular cell biology 11(2-3): 377–394.

Yoh SM, Lucas JS, Jones KA. 2008. The Iws1:Spt6:CTD complex controls cotranscriptional mRNA biosynthesis and HYPB/Setd2-mediated histone H3K36 methylation. Genes & development 22(24): 3422–3434.

Yuan W, Xie J, Long C, Erdjument-Bromage H, Ding X, Zheng Y, Tempst P, Chen S, Zhu B, Reinberg D. 2009. Heterogeneous nuclear ribonucleoprotein L Is a subunit of human KMT3a/Set2 complex required for H3 Lys-36 trimethylation activity in vivo. The Journal of biological chemistry 284(23): 15701–15707.

Zhang D, Yoon HG, Wong J. 2005. JMJD2A is a novel N-CoR-interacting protein and is involved in repression of the human transcription factor achaete scute-like homologue 2 (ASCL2/Hash2). Molecular and cellular biology 25(15): 6404–6414.

Zhou HL, Luo G, Wise JA, Lou H. 2014. Regulation of alternative splicing by local histone modifications: potential roles for RNA-guided mechanisms. Nucleic acids research 42(2): 701–713.

